# Endothelial alpha globin is a nitrite reductase

**DOI:** 10.1101/2021.02.12.430954

**Authors:** T.C. Stevenson Keller, Alexander S. Keller, Gilson Brás Broseghini-Filho, Joshua T. Butcher, Henry R. Askew Page, Aditi Islam, Zhe Yin Tan, Leon J. DeLalio, Christophe Lechauve, Steven Brooks, Poonam Sharma, Kwangseok Hong, Wenhao Xu, Alessandra Simão Padilha, Claire A. Ruddiman, Angela K. Best, Edgar Macal, Daniel B. Kim-Shapiro, George Christ, Zhen Yan, Miriam M. Cortese-Krott, Karina Ricart, Rakesh Patel, Timothy P. Bender, Swapnil K. Sonkusare, Mitchell J. Weiss, Hans Ackerman, Linda Columbus, Brant E. Isakson

## Abstract

Small artery vasodilation in response to hypoxia is essential for matching oxygen supply to tissue oxygen demand. One source of hypoxic dilation via nitric oxide (NO) signaling is nitrite reduction by erythrocytic hemoglobin (α2β2). However, the alpha subunit of hemoglobin is also expressed in resistance artery endothelium and localized to myoendothelial junctions, a subcellular domain that contacts underlying vascular smooth muscle cells. We hypothesized that nitrite reduction mediated by endothelial alpha globin may occur at myoendothelial junctions to regulate hypoxic vasodilation. To test this concept, we created two novel mouse strains: one lacking alpha globin specifically in endothelium (EC *Hba1*^*Δ/Δ*^) and one where alpha globin is mutated such that its inhibitory association with endothelial NO synthase (eNOS) is prevented (*Hba1*^*WT/Δ36-39*^). In EC *Hba1*^*Δ/Δ*^ or *Hba1*^*WT/Δ36-39*^ mice hemoglobin levels, hematocrit and erythrocyte counts were unchanged from littermate controls. Loss of the full alpha globin protein from the endothelium in the EC Hba1^Δ/Δ^ model was associated with decreased exercise capacity and decreased intracellular nitrite utilization in hypoxic conditions. These effects were not seen in *Hba1*^*WT/Δ36-39*^ animals. Hypoxia induced vasodilation was decreased by 60% in isolated thoracodorsal arteries from EC *Hba1*^*Δ/Δ*^, while infusion of erythrocytes only partially rescued the dilatory response. Lastly, unlike other models where blood pressure is decreased, EC *Hba1*^*Δ/Δ*^ blood pressure was not altered in response to hypoxia. Overall, we conclude that alpha globin in the resistance artery endothelium can act as a nitrite reductase to provide a local vasodilatory response to hypoxia.

## Introduction

Matching blood perfusion to metabolic demand is a critical function of the autoregulation of the resistance vasculature. Although a number of vasodilatory signaling pathways have been identified in small arteries, nitric oxide (NO) is among the most potent mechanisms. This is especially relevant in hypoxia, where NO has been implicated as a key regulator of acute hypoxic vasodilatory responses^1, 2, 3^.

A major source of hypoxic NO generation is the reaction of the nitrite anion with metal centers, especially the iron in erythrocytic hemoglobin ^2, 4^. This reaction occurs with deoxygenated heme groups, where nitrite is reduced to vasoactive NO. Many studies from independent laboratories have implicated erythrocyte NO generation and/or delivery as a critical source of hypoxic vasodilation, even as some questions remain about the nature and stability of the molecule carrying “NO bioactivity” ^5, 6, 7, 8^. Other sources of NO for hypoxia-driven vasodilation in the resistance vasculature have yet to be determined.

Our group has previously described alpha globin in the endothelium of resistance arteries localized specifically at the site of endothelial and smooth muscle interaction, the myoendothelial junction (MEJ)^9, 10, 11^. Here, the alpha globin acts as a negative regulator of endothelial NO synthase (eNOS) derived signaling through NO scavenging aided by direct interaction with eNOS^9, 10, 12, 13^. Negative regulation of NO signaling by endothelial alpha globin is O_2_ dependent, as both eNOS production of NO and deactivation of NO by alpha globin (producing nitrate, NO_3_^−^) require O_2_. Recent work from human studies confirmed an role for endothelial alpha globin in NO catabolism, as alpha thalassemic individuals demonstrated increased flow-mediated dilation^14^.

Endothelial alpha globin is also a candidate for controlling hypoxia-induced vasodilation by generating NO through nitrite reduction. Expression of alpha globin in the MEJ presents a pool of hemoproteins that are optimally positioned next to vascular smooth muscle to control vasodilation. Endothelial alpha globin might act as a sensor and actuator of hypoxia-driven signaling, effecting precise control of local vasodilation to perfuse metabolically active tissues during exercise or other acute hypoxia scenarios. We hypothesized that endothelial alpha globin has a critical secondary role in controlling vasodilation separate from its catabolic role in NO homeostasis through interaction with eNOS.

To test whether endothelial alpha globin contributes to hypoxic vasodilation signaling, we have generated two genetic mouse models: one with endothelial-specific loss of alpha globin through *Cre/lox* recombination and a CRISPR-based mutant in which alpha globin is present but lacks its eNOS-binding domain. Using these two models, we have decoupled the function of alpha globin as an eNOS-interacting NO scavenger from its proposed role as a nitrite reductase. We further show deficiencies in hypoxia-induced vasodilation, as well as global physiologic measurements of exercise capacity and blood pressure regulation only when the full alpha globin protein is deleted. Overall, we demonstrate that endothelial alpha globin can act as a nitrite reductase further refining the roles of alpha globin in regulating vascular NO homeostasis.

## Methods

### Animals

*Hba1*^*fl/fl*^ mice were developed in partnership with Ingenious Targeting Laboratory. *Hba1*^*fl/fl*^ mice were crossed with the *Cdh5-PAC-Cre*^*ERT2*^ transgenic driver to yield a model of tamoxifen-inducible, endothelial-cell-specific *Hba1* deletion. EC Hba1^Δ/Δ^ mice and Cre (-) littermate controls received 10 days of tamoxifen injections (100 μl at 10 μg/ml) starting at 6 weeks of age. Both males and females were used for experiments. Mice were maintained on Teklad irradiated rodent diet (Teklad product number 7912). All animals were used between 12 and 20 weeks of age. All animals were handled according to protocols approved by the University of Virginia IACUC.

### Generation of CRISPR/Cas9 constructs

*HBA-a1* (NC_000077.6) targeting site for guide RNAs were identified using the online Zifit Targeter software version 4.2 (http://zifit.partners.org)^15^. The sgRNA target site 5’- GGAAAGTAGGTCTTGGTGGT −3’ and its recommended 22 nucleotide long oligos(oligo 1: 5’- TAGGAAAGTAGGTCTTGGTGGT −3’; oligo 2: 5’- AAACACCACCAAGACCTACTTT −3’) were chosen from Zifit output. Oligo1 and oligo 2 were annealed and cloned at BsaI site in DR274 vector (http://www.addgene.org/crispr/jounglab/crisprzebrafish/) followed by transformation in chemically competent XL-1 Blue cells. Antibiotic resistant positive clones were checked by sequencing with M13 primers. Positive PDR274 clones were digested with Dra1 restriction enzyme (New England Bioscience) and used as a template for in vitro transcription using MEGAshortscript T7 Transcritption Kit (Life Technologies). sgRNAs were DNAse treated and purified using the MEGAclear Kit (Ambion P/N AM1908). Purity and concentration were measured using a NanoDrop spectrophotometer.

Production of Cas9 nuclease mRNA utilized the pMLM313 expression vector (Addgene plasmid #42251), which harbors the T7 promoter site upstream of the translational start site, and a nuclear localization signal at the carboxy-terminal. The pMLM313 plasmid was linearized using Pme1 restriction enzyme (New England Bioscience), separated by agarose gel, and gel extracted (Clontech Nucleospin Extract II Kit Cat# 636972). The linearized and purified pMLM313 fragment was used as template for *in vitro* transcription of Cas9 mRNA with mMESSAGE mMACHINE T7 ULTRA kit (Life Technologies)^16, 17, 18^. Following transcription, Cas9 mRNA underwent poly-A tailing reactions and DNase I treatment according to manufacturer’s instructions. The concentration was determined using NanoDrop.

### Microinjection

Pronuclei of one-cell stage embryos were collected from superovulated donor females (B6/SJL strain) and microinjected with CRISPR/Cas9 constructs using micromanipulators and injection needles (World Precision Instruments, filament #1B100-6). Injected embryos were implanted into pseudopregnant recipient females according to NIH guidelines. 2-week-old pups born to recipient females were screened by PCR using *Hba1* gene specific primers and sequenced for the presence of CRISPR/Cas9 induced mutation. The *Hba1*-gen-FP primers were forward: 5’- ATATGGACCTGGCACTCGCT −3’ and reverse: 5’- GTCCCAGCGCATACCTTGAA −3’.

### Whole genome sequencing

Whole genome sequencing was performed by GeneWiz (South Plainfield, NJ) using contiguous amplicon sequencing on Illumina MiSeq instrument in a 2×300bp paired-end configuration. DNA library preparation without fragmentation, multiplexing, sequencing, unique sequence identification, and relative abundance calculations were performed in house at GeneWiz.

### Blood cell measurements

Isolated mouse blood was analyzed by the St. Jude Children’s Hospital Blood Pathology Labs as previously described by us^9^.

### Immunofluorescence

Thoracodorsal arteries were excised, cleared of blood, and fixed overnight in 4% paraformaldehyde. After washing with ethanol, vessels were embedded in agarose and then paraffin blocks, and sectioned at 5 μm. Paraffin was removed by Histoclear, and sections were rehydrated through a 100%/95%/70% ethanol/water gradient before antigen retrieval in a citrate solution. PBS blocking solution (including 0.25% Triton X-100, 0.5% bovine serum albumin, and 5% donkey serum) was used to block sections and as primary and secondary diluent. Primary antibodies were rabbit anti-hemoglobin alpha (abcam, #102758, 1:250 dilution) and mouse anti-eNOS (BD Bioscience, #610296, 1:500 dilution). Secondary antibodies used were donkey anti-rabbit (Alexa 647, Life Technologies A-31571) and donkey anti-mouse (Alexa 594, Life Technologies A-21207). Stained sections were mounted with DAPI and imaged using an Olympus FV1000 confocal microscope with Olympus Fluoview software using DAPI, and Alexa 488 (to visualize autofluorescence of the elastic lamina), 594, and 647 channels for fluorescence detection.

### Detection of globin precipitates from RBCs

Analysis of globin precipitates from erythrocytes was performed as described.^19, 20, 21, 22, 23^ Briefly, washed RBCs were lysed and pellets washed extensively in ice-cold 0.05X PBS. Membrane lipids were extracted with 56 mM sodium borate, pH 8.0 with 0.1% Tween-20 at 4°C. Precipitated globins were dissolved in 8M urea, 10% acetic acid, 10% b-mercaptoethanol, and 0.04% pyronin, fractionated by triton-acetic acid-urea (TAU) gel electrophoresis and stained with Coomassie blue.

### Fluorescence polarization

Fluorescently labeled peptides were synthesized by *Anaspec*. The fluorescence polarization was performed as described previously by us^9^. Briefly, 1 μM recombinant oxygenase domain of eNOS was incubated with a peptide composed of the alpha globin binding sequence (LSFPTTKTYF) or the same peptide lacking *Hba1* amino acids 36-39 (sequence: LTKTYF) to assess binding affinity. Each peptide was synthesized with an Alexa Flour 488 (excitation 494 nm, fluorescent lifetime 4.1 ns) tag at the C-terminus to perform this assay.

### Proximity ligation assay

Proximity ligation assays (PLA) were performed as described by us^11^. Sections (as in “Immunofluorescence”) were stained with the same primary antibodies at the same concentrations. DUOLINK® PLA probes, ligase, and polymerase were all used according to manufacturer’s instructions. Sections were mounted with DAPI, and imaged with DAPI, Alexa 488, and Texas Red (for the PLA fluorophore) channels. Images of two sections per animal (with genotype blinded) were quantified by tracing the outline of the IEL in ImageJ and dividing number of punctae by total perimeter of elastic lamina.

### Immunoblots

Immunoblots were quantified against total protein using a Licor Odyssey with near-infrared fluorescent secondary antibodies as previously described by us^24^.

### Nitric oxide imaging

Arterioles were surgically opened and pinned down on a Sylgard block in *en face* preparation. NO levels were assessed using 5 μM DAF-FM DA (4-amino-5 methylamino-2’,7’-difluorofluorescenin diacetate) prepared in HEPES-PSS (physiologic saline solution) with 0.02% pluronic acid.^25^ DAF-FM DA reacts with NO in the presence of oxidants and forms a fluorescent triazole compound. *En face* mesenteric arteries were pre-treated with carbachol (CCh, muscarinic receptor agonist, 10 μM) in Hepes-PSS for 5 minutes at 30°C. Arteries were then incubated with DAF-FM containing CCh for 20 minutes at 30°C in the dark. DAF-FM DA fluorescence was imaged using Andor Revolution WD (with Borealis) spinning-disk confocal imaging system (Andor Technology, Belfast, UK) comprised of an upright Nikon microscope with a water dipping objective (60X, numerical aperture (NA) 1.0) and an electron multiplying CCD camera. Images were obtained along the z-axis at a slice size of 0.05 μm from the top of the endothelial cells to the bottom of the smooth muscle cells using an excitation wavelength of 488 nm and emitted fluorescence was captured with a 525/36-nm band-pass filter. DAF-FM fluorescence was analyzed using custom-designed SparkAn software by Dr. Adrian D. Bonev (University of Vermont, Burlington, VT USA). An outline was drawn around each endothelial or smooth muscle cell to obtain the arbitrary fluorescence intensity of that cell. The background fluorescence was then subtracted from the recorded fluorescence. The fluorescence from all the cells in a field of view was averaged to obtain single fluorescence number for that field. Relative changes in DAF-FM fluorescence were obtained by dividing the fluorescence in the treatment group by that in the control group. Each field of view was considered as n=1, and several fields of view from at least three arteries from at least three mice from each group were included in the final analysis.

### Treadmill running test

Mice aged 10-13 weeks were acclimatized to the treadmill in accordance with established methods^26^. On the day of testing, the treadmill was set to 5% incline at 13 m/min (0.5 miles per hour) for 30 minutes, and increased by 2.7 m/min (0.1 miles per hour) every 30 minutes (to a maximum of 27 m/min [1.0 miles per hour]) until mice reached exhaustion. Exhaustion was defined as refusal to run when induced by 20 strokes on the tail with a wire brush. Blood lactate was measured immediately before and after the test with tail snip and lactate measurement using a Lactate Scout handheld monitor. All mice were monitored by an investigator blinded to genotypes.

### Nitrite/Nitrate Measurement

Vessels were blotted dry, and 50ul PBS containing 2.5mM EDTA, 100mM NEM, pH7.4 added. Vessels were sonicated using a Sonic dismembrator model 100 (Fisherbrand), twice, 10 sec each cycle (0.05 watts). Then, methanol was added (100 μl) and samples vortex mixed for 20 sec followed by centrifugation (19000 x g, 10min at 4°C). 100 μl of the supernatant was taken and nitrite and nitrate measured using an NOX analyzer (ENO-30). Concentrations were calculated by comparison to freshly prepared sodium nitrite and sodium nitrate standard curves.

### Vasoreactivity

Cumulative dose-response curves on pressurized and cannulated thoracodorsal arteries (140-200 μm in diameter) were used throughout the studies as previously described. Briefly, freshly isolated thoracodorsal arteries were placed into ice-cold Krebs-HEPES solution containing (in mM) NaCl 118.4 KCl 4.7, MgSO_4_ 1.2, NaHCO_3_ 4, KH_2_PO_4_ 1.2, CaCl_2_ 2, HEPES 10, and Glucose 6. The vessels were then mounted in a pressure arteriograph (Danish MyoTechnology) with the lumen filled with Krebs-HEPES solution. The vessels were pressurized to 80 mmHg and heated to 37°C. Between experiments, bath solution was washed out for 10 minutes with fresh Krebs-HEPES buffer, after which a new Krebs-HEPES solution was added and allowed to re-equilibrate. All vessels were tested for endothelial function before NaS_2_O_4_experiments by assessing dilatory response to 1 μM NS309. Phenylephrine preconstriction diameter was determined by addition of 1 μM phenylephrine to the bath, and waiting until a stable plateau of internal diameter, before continuing to the next stimulation condition. Following conclusion of experimental stimulations, maximal constriction of vessels was determined using a stimulation of 3 0mM KCl, and finally vessels were washed with a Ca^2+^-free Krebs-HEPES solution supplemented with 1 mM EGTA and 10 μM sodium nitroprusside to determine maximal passive diameter. Dilation to NaS_2_O_4_ was calculated as (diameter after NaS_2_O_4_ stimulation – diameter before NaS_2_O_4_ stimulation) / (passive diameter – diameter before NaS_2_O_4_ stimulation) * 100.

### Blood pressure

Radiotelemetry was used to monitor blood pressure and heart rate responses in conscious animals, as previously described by us^9^.

### Statistics

In all cases, mean values are shown with error bars denoting SEM unless otherwise stated. Statistical tests are denoted in the figure captions below each figure. Graphpad Prism 9 was used to analyze statistical relationships between groups.

## Results

### Alpha globin is the only globin expressed in the endothelium of resistance vasculature

Expression of the alpha subunit of hemoglobin has been observed in the endothelium of small resistance arteries by our group and others^10, 11, 12, 27^. Before targeting alpha globin, we assayed for other mammalian globins (including myoglobin, cytoglobin, and neuroglobin) that could catalyze nitrite reduction in the vascular wall of resistance arteries. Thus, using the murine skeletal muscle-supplying thoracodorsal artery as a model of resistance arteries (an artery with average diameter less than 200 μm), and murine aorta as a representative conduit artery, we assayed expression of globin isoforms in endothelium and smooth muscle layers using confocal microscopy (**Figure 1**). Human small arteries from adipose biopsies were also used as a translational comparison (**Supplemental Figure 1**). In the murine thoracodorsal artery, a strong signal for alpha globin is observed in the endothelium (**Figure 1A**, vessel lumen denoted by an asterisk). No other globin isoforms are observed in the endothelium, though cytoglobin and neuroglobin appear localized to adventitial layers in this artery (**Figure 1A**). Similarly, alpha globin was the only endothelial globin observed in the tested human microvessels. Weak staining was observed for cytoglobin, myoglobin, and neuroglobin in the smooth muscle and adventitial layers (**Supplemental Figure 1A**). Alpha globin expression in the endothelium is characteristic of resistance vessels and is not found in all vessel types; in the aorta, alpha globin is absent from the endothelium (**Figure 1B**). A strong myoglobin signal was observed in the multiple smooth muscle layers of the aorta (**Figure 1B**). Antibody isotype controls were analyzed (**Supplemental Figure 1B**). Positive immunofluorescence signal for hemoglobin β was observed only in erythrocytes (**Supplemental Figure 2**).

**FIGURE 1:**
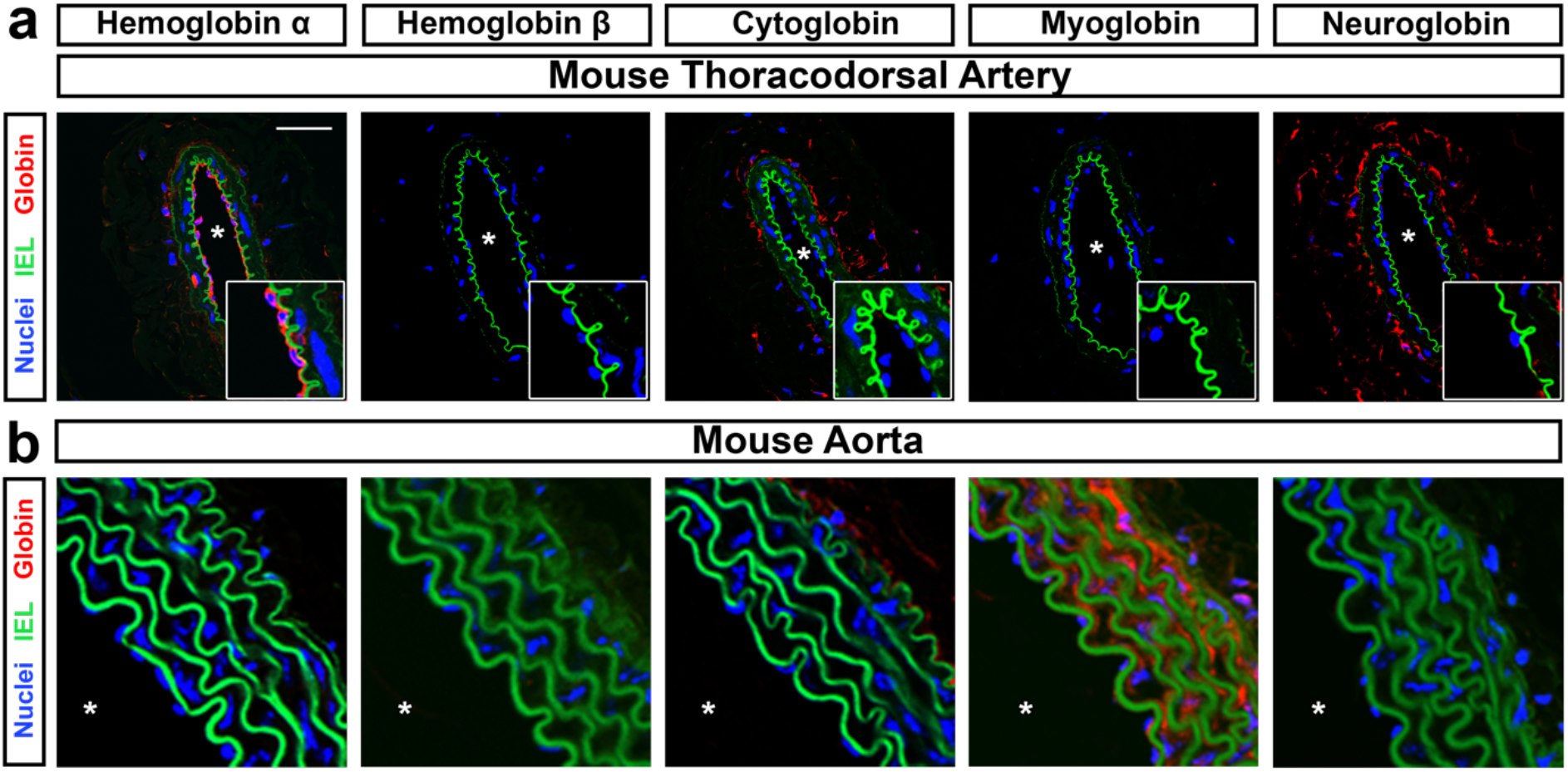
Alpha globin is the only globin expressed in the mouse endothelium. (**a**) The mouse thoracodorsal artery (a skeletal muscle-derived resistance artery) shows endothelial expression of alpha globin, but not other globins. Cytoglobin and neuroglobin are observed in adventitial layers, but not endothelial cells. (**b**) The mouse aorta is representative of globin expression in a conduit artery. Only myoglobin shows expression in the aorta wall, but not endothelium. The scale bar in A represents 50 μm; zoomed images in focused on the endothelium are shown in the bottom right. In all images, the asterisk denotes the lumen of the vessel. All images are representative of observations are from a minimum of 4 mice per staining condition.

### Generation of an Hba1 conditional allele

To test the role of alpha globin in endothelium, we generated a mutant *Hba1* allele with *loxP* recombination sites flanking exons 2 and 3 (**Figure 2A**). These mutant mice, bred to homozygosity (*Hba1*^*fl/fl*^), present with no observed lethality (**Supplementary Table 1**). Temporal and endothelial cell type-specific knockout of *Hba1* was driven by the tamoxifen-inducible Cre recombinase downstream of a VE-Cadherin promoter (*Cdh5-PAC-Cre*^*ERT2*^)^28^ (denoted EC *Hba1*^*Δ/Δ*^ hereafter) (**Figure 2A**). After recombination, PCR amplification of the *Hba1* locus demonstrates a smaller amplification product (450 bp compared to 1100 bp in control animals) in EC *Hba1*^*Δ/Δ*^ mice due to deletion of exons 2 and 3 (**Figure 2B**) with no change in expected birth ratios with the Cre driver (**Supplemental Table 2**). Endothelial-specific knockout of *Hba1* reduces immunofluorescent signal of alpha globin in the endothelium, in particular in holes in the internal elastic lamina where myoendothelial junctions form, compared to littermate controls (**Figure 2C-E**). We do not observe difference in blood counts in EC *Hba1*^*Δ/Δ*^, as would be expected from loss of *Hba1* in cells of myeloid lineages (**Figure 2F-H**). There were no changes in erythrocyte size or volume upon *Cdh5-Cre* driven deletion (**Supplementary Figure 3**); this is in contrast to the anemia and decreased red blood cell size in *Hba1* global knockout animals (*Hba1*^*-/-*^, **Supplemental Figures 3-4**). Global recombination of our *Hba1* conditional allele (*Hba1*^*fl/fl*^; *Sox2-Cre*^29^), hereafter *Hba1*^*Δ/Δ*^, **Supplemental Figure 4A-C**) produces an anemia phenotype that mirrors global loss of *Hba1* through insertion of a neomycin resistance cassette^30^ (**Supplemental Figure 4D-F**) with significant defects in blood hemoglobin and hematocrit without a statistically significant change in total red blood cells.

**FIGURE 2:**
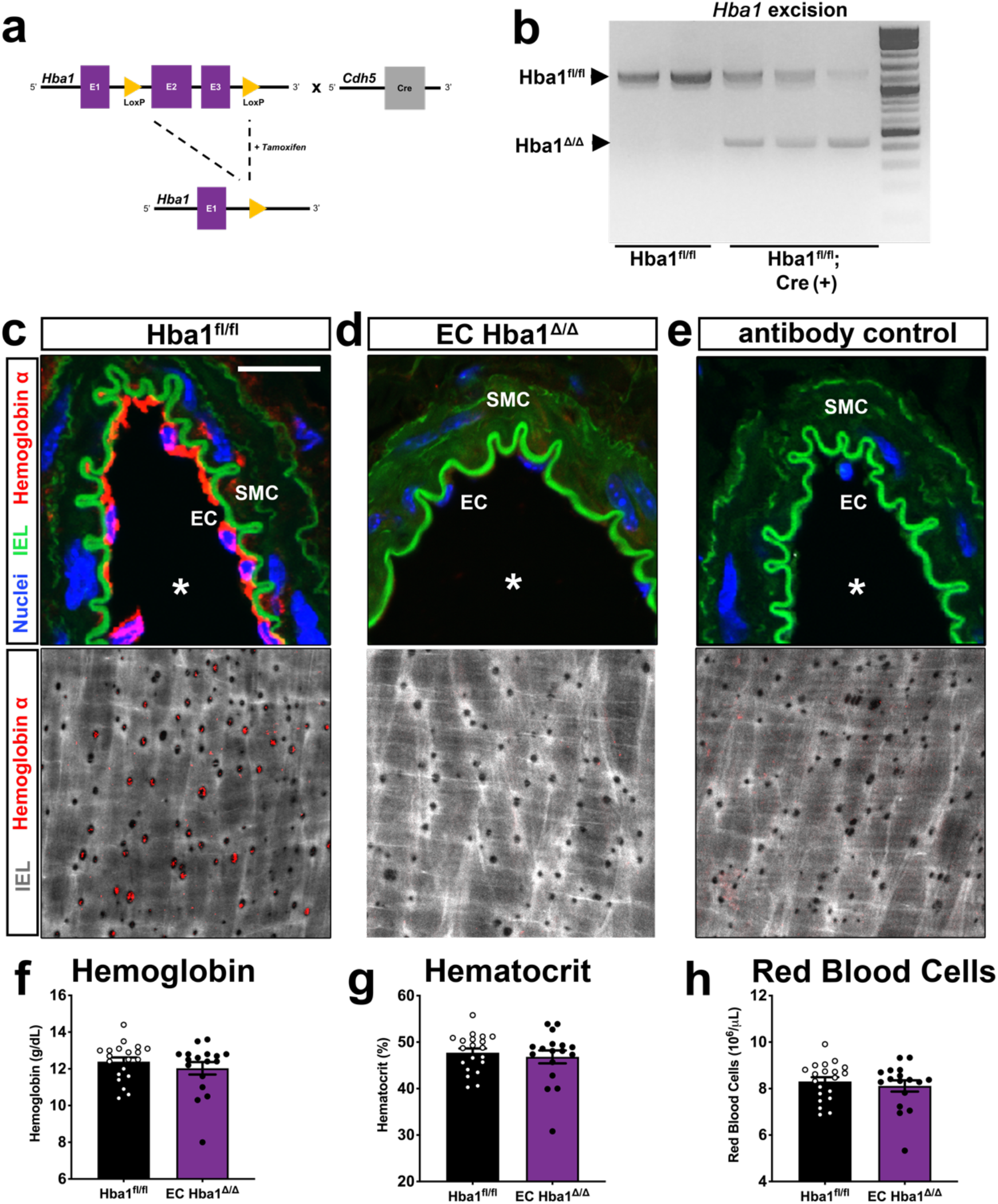
Creation of an endothelial-specific Hba1 deletion mouse model. (**a**) Deletion strategy: *loxP* sites flanking exons 2 and 3 of the mouse *Hba1* gene (*Hba1*^*fl/fl*^) were introduced by recombineering. A tamoxifen-inducible, endothelial-specific Cre recombinase (*Cdh5-PAC-Cre*^*ERT*^2^^) allows for temporally controlled and cell type-specific deletion of a functional *Hba1* gene (EC *Hba1*^*Δ/Δ*^ mouse). (**b**) DNA gel showing recombination of the *Hba1* locus with Cre activation. The recombination event produces a band at ∼450 bp. (**c**, **d**) Immunofluorescence staining for alpha globin in transverse sections (top; thoracodorsal artery) and *en face* views (bottom; third order mesenteric artery). Scale bars in both are 25 μm and * indicates lumen of the vessel in top images. Alpha globin (red) was found in the endothelium throughout, and specifically in the holes in the internal elastic lamina (IEL) where myoendothelial junctions are found. (**e**) Control IgG staining shows the specificity of the staining for alpha globin in this tissue. (**f-g**) endothelial deletion of alpha globin does not affect blood cell hemoglobin parameters. Blood hemoglobin content (**f**), hematocrit (**g**), and the number of red blood cells (**h**) is unchanged in the EC *Hba1*^*Δ/Δ*^ mouse compared to the *Hba1*^*fl/fl*^ control. For experiments in f-h, n = 20 *Hba1*^*fl/fl*^ and n = 17 EC *Hba1*^*Δ/Δ*^; t-tests were used to determine if a significant difference exists between groups.

Further, the specificity of the recombination event for the *Hba1* gene locus is demonstrated by the deletion with the *Sox2-Cre*. If both *Hba1* and *Hba2* were affected by the *Cre/lox* recombination, severe alpha thalassemia would be induced, resulting in embryonic lethality^31^. *Hba1*^*Δ/Δ*^ mice are born at expected ratios (**Supplemental Table 3**, compare with genotypes from the global *Hba1*^*-/-*^ model in **Supplemental Table 4**), suggesting that only the *Hba1* locus is flanked by recombination sites, even as *Hba1* and *Hba2* loci share high sequence similarity.

### Generating a genetic model to disrupt alpha globin/eNOS interaction

Endothelial specific-knockdown of the total alpha globin protein prevents interrogation of our hypothesized nitrite reduction role independent of its known role in association with eNOS^9,10^. To uncouple the effects of loss of the heme site and eNOS binding, we generated a murine model where alpha globin is specifically lacking the identified eNOS-interacting sequence (*Hba1* residues 34-43) ^9, 10^. Using CRISPR/Cas9 gene editing to target the region of alpha globin known to interact with eNOS (**Figure 3A**), a genetic model was generated with a mutation (in-frame deletion of 12 nucleotides, **Figure 3B and C**) that removed *Hba1* residues 36-39 (NH_2_ – SFPT – COOH). We refer to this allele as *Hba1*^*Δ36-39*^. The expression of the mutant alpha globin protein was observed in immunoprecipitate from lysed red blood cells, where an approximate 400 Da peak shift (corresponding to loss of 4 amino acids from the polypeptide chain) was observed in a matrix-assisted laser desorption ionization mass spectrometer (**Figure 3D**).

**FIGURE 3:**
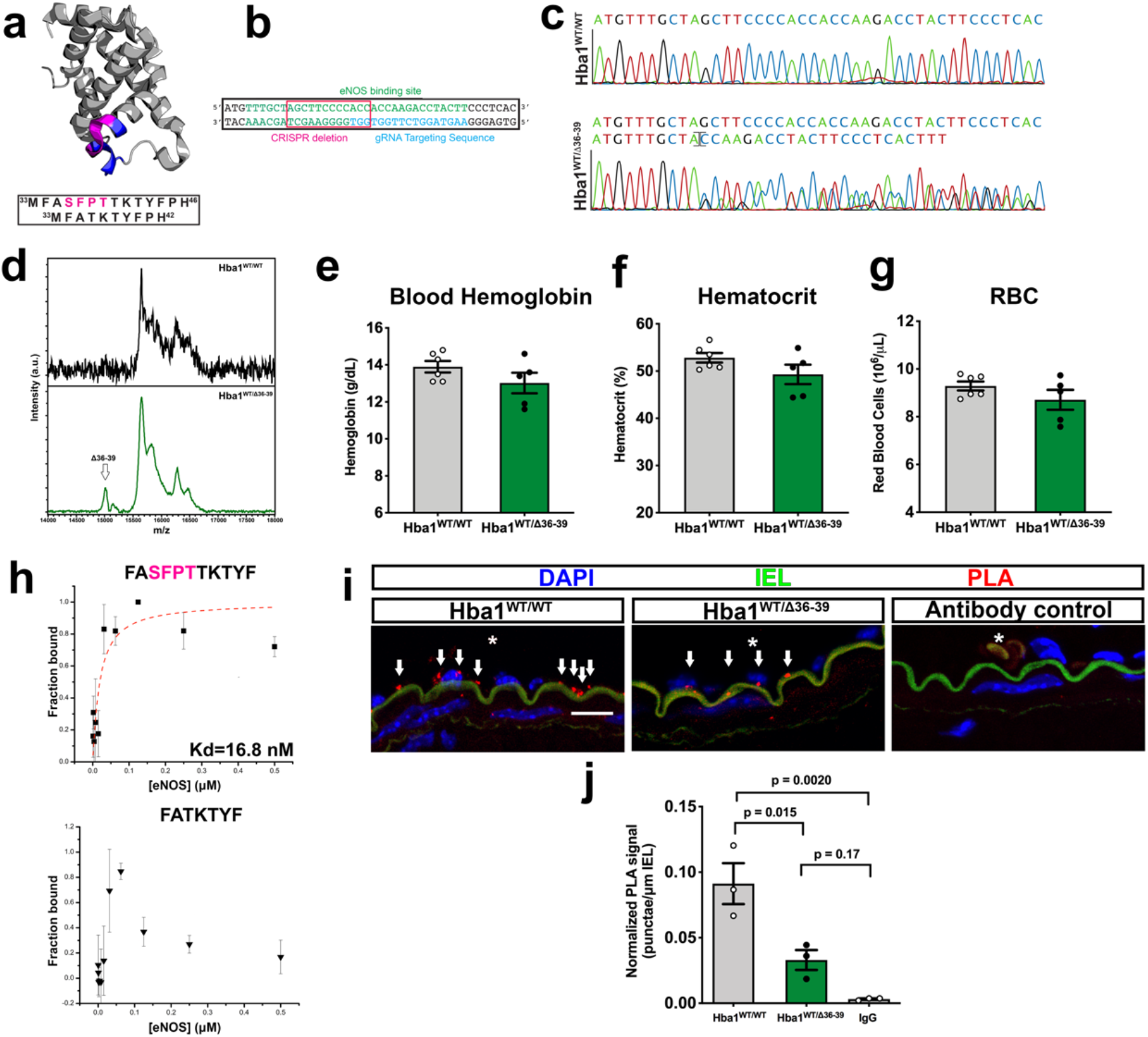
Disruption of alpha globin/eNOS interaction through deletion of four residues in Hba1. A colorized crystallographic model of alpha globin (from PDBid: 1Z8U) showing the residues previously shown to interact with eNOS (blue) surrounding four residues deleted in the *Hba1*^*WT/Δ^36-39^*^ mouse model (magenta). The sequences for the *Hba1*^*WT*^ and *Hba1*^*Δ*36-39^ alleles, including the deleted residues in magenta, are shown in the box below. (**b**) A guide RNA design (blue) targeting the eNOS binding region (green) resulted in an in-frame deletion of the nucleotides boxed in pink. This is confirmed by the chromatogram in (**c**), where the twelve nucleotide deletion scrambles downstream reading of the nucleotide sequence in NGS protocols. (**d**) The mutant protein is expressed, as seen in immunoprecipitation-coupled mass spectrometry. A peak corresponding to a protein molecular weight ∼400 Da less than the dominant species is seen in hemoglobin captured from lysed red blood cells. (**e-g**) Blood hemoglobin content (**e**), hematocrit (**f**), and number of red blood cells (**g**) are shown from *Hba1*^*WT/WT*^ and *Hba1*^*WT/Δ^36-39^*^ groups. (**h**) Fluorescence polarization assays for determining the binding of the mutant allele to eNOS, using an alpha globin mimetic peptide known to bind (top), compared to the Δ36-39 peptide on the bottom. No binding affinity can be calculated for the Δ36-39 peptide. Points are centered on the mean value of 3 measurements per concentration, and error bars represent standard deviation of the triplicate measurements. (**i**) Proximity ligation assay (PLA) signal (red punctae, marked by arrows) highlights close localization of alpha globin and eNOS. Nuclei are seen in blue (DAPI), the autofluorescence of the internal elastic lamina in green, and the lumen of the vessel is denoted by an asterisk (*). Scale bar is 15 mm. (**j**) Endothelial PLA signals, normalized to length of the internal elastic lamina, are reduced in the *Hba1*^*WT/Δ^36-39^*^ mice (significantly different from *Hba1*^*WT/WT*^, but not from IgG staining controls). For **f-h**, n = 6 for *Hba1*^*WT/WT*^ and n = 5 for *Hba1*^*WT/Δ^36-39^*^. For **j**, n = 3 in each genotype, and differences were determined by a t-test with multiple comparisons correction.

The global deletion of these four residues was embryonically lethal by E14.5 in homozygosity (*Hba1*^*Δ36-39/Δ36-39*^; **Supplemental Table 5**), showing growth restriction in gross observation, and dilated major vessels and eccentrically dilated, non-compact nascent myocardium via histology (**Supplemental Figure 5A-B**). A large increase in tyrosine nitration (via immunofluorescence staining) is a possible downstream effect of oxidative and nitrosative stress due to high output NO generation during development (**Supplemental Figure 5C**). Homozygous mutants were not achieved in sufficient numbers for experimentation, so we used heterozygous *Hba1*^*WT/Δ36-39*^ mice which were fertile and viable into adulthood (**Supplemental Table 5**).

The *Hba1*^*WT/Δ36-39*^ mice were hypothesized to have decreased alpha globin/eNOS interaction, allowing increased NO availability in endothelium. In line with increased tyrosine nitration observed in the *Hba1*^*Δ36-39/Δ36-39*^ embryos, adult *Hba1*^*WT/Δ36-39*^ mice had a significantly increased NO production after cholinergic stimulation (**Supplemental Figure 6**), which would be predicted based on previous results with an exogenous peptide (HbαX) that we have shown disrupts alpha globin/eNOS complex formation^9, 10^. Thus, we were able to create a mouse model that mimicked the effects of eNOS and alpha globin loss of interaction (as in prior work with HbαX), but retains the alpha globin in the endothelium.

First, we determined whether the *Hba1*^*Δ36-39*^ mutation affects erythrocytes in adult heterozygous mice. *Hba1*^*WT/Δ36-39*^ mice did not present with abnormalities in total blood hemoglobin, hematocrit, or number of red cells (**Figure 3E-G**). Further, there were no observed differences in mean corpuscular volume or mean corpuscular hemoglobin for red cells (**Supplemental Figure 7A-B**). Erythrocytes from the *Hba1*^*WT/Δ36-39*^ mouse did not appear to turn over more rapidly as the percentage of reticulocytes was unchanged (**Supplemental Figure 7C**). Hemoglobin tetramers from the *Hba1*^*WT/Δ36-39*^ mice were stable, as no insoluble alpha globin was found in blood samples isolated from these mice (**Supplemental Figure 7D**). Additionally, red blood cell morphology is normal with no indication of increased hemoglobin precipitation within the red cell stained by Cresyl Blue (**Supplemental Figure 7E, F**).

To determine that deletion of ^36^SFPT^39^ in alpha globin prevents alpha globin/eNOS interaction in the endothelium, *in vitro* studies of binding affinity were performed (**Figure 3H**). A peptide corresponding to the region of alpha globin that binds with eNOS (*Hba1* residues FASFPTTTKTYF, **Figure 3H, top**) bound to the recombinant eNOS oxygenase domain with an affinity similar to previous studies (16.8 nM)^9^. In contrast, the Δ36-39 deletion peptide (**Figure 3H, bottom**) did not bind to the eNOS oxygenase domain with a detectable affinity using this methodology. The loss of the interaction was assayed in *ex vivo* vessels using a proximity ligation assay (PLA). Using this technique, interaction of alpha globin and eNOS was found to be significantly reduced in endothelium of the *Hba1*^*WT/Δ36-39*^ mouse (**Figure 3I-J**). Arteries from *Hba1*^*WT/Δ36-39*^ and *Hba1*^*WT/WT*^ did not have observable difference in alpha globin and eNOS expression in endothelium (**Supplemental Figure 8**).

### Treadmill running distance is moderately decreased in EC Hba1^Δ/Δ^ mice

A major physiologic role for hypoxic vasodilation is regulating blood perfusion to match metabolic demand. This is particularly relevant in the context of exercise-induced hyperemia in skeletal muscles. A forced exercise model can recapitulate conditions of relative hypoxia in skeletal muscle^32, 33^; we used this experiment to determine whether loss of putative nitrite reductase function from knockout of endothelial alpha globin decreases vasodilation to hypoxic stress and thus exercise fitness. Mice were encouraged to run on a treadmill with controlled speed until exhaustion (**Figure 4A**). Between littermate controls of the EC *Hba1*^*Δ/Δ*^ and *Hba1*^*fl/fl*^ mice, as well as the *Hba1*^*WT/WT*^ and *Hba1*^*WT/Δ36-39*^ mice, there were no differences in body weight (**Figure 4B**). Distance to exhaustion was significantly decreased in the EC *Hba1*^*Δ/Δ*^ group compared to *Hba1*^*fl/fl*^ genotypes. *Hba1*^*WT/WT*^ and *Hba1*^*WT/Δ36-39*^ groups were exhausted at a similar distance (**Figure 4C**). Blood lactate measurements (a metric for muscular workload) were not significantly different between littermate-paired groups (**Figure 4D**). Similar to EC *Hba1*^*Δ/Δ*^, global knockout of *Hba1* (in the *Hba1*^*Δ/Δ*^) model caused a significant decrease in running distance (**Supplemental Figure 9**). The effects of mouse genotype on running distance were independent of differences in soleus muscle capillary density (**Figure 4E**). Taken together, it appears that preventing alpha globin/eNOS association does not affect exercise capacity. However, loss of endothelial alpha globin protein, including possible nitrite reduction capacity, is detrimental to exercise capacity.

**FIGURE 4:**
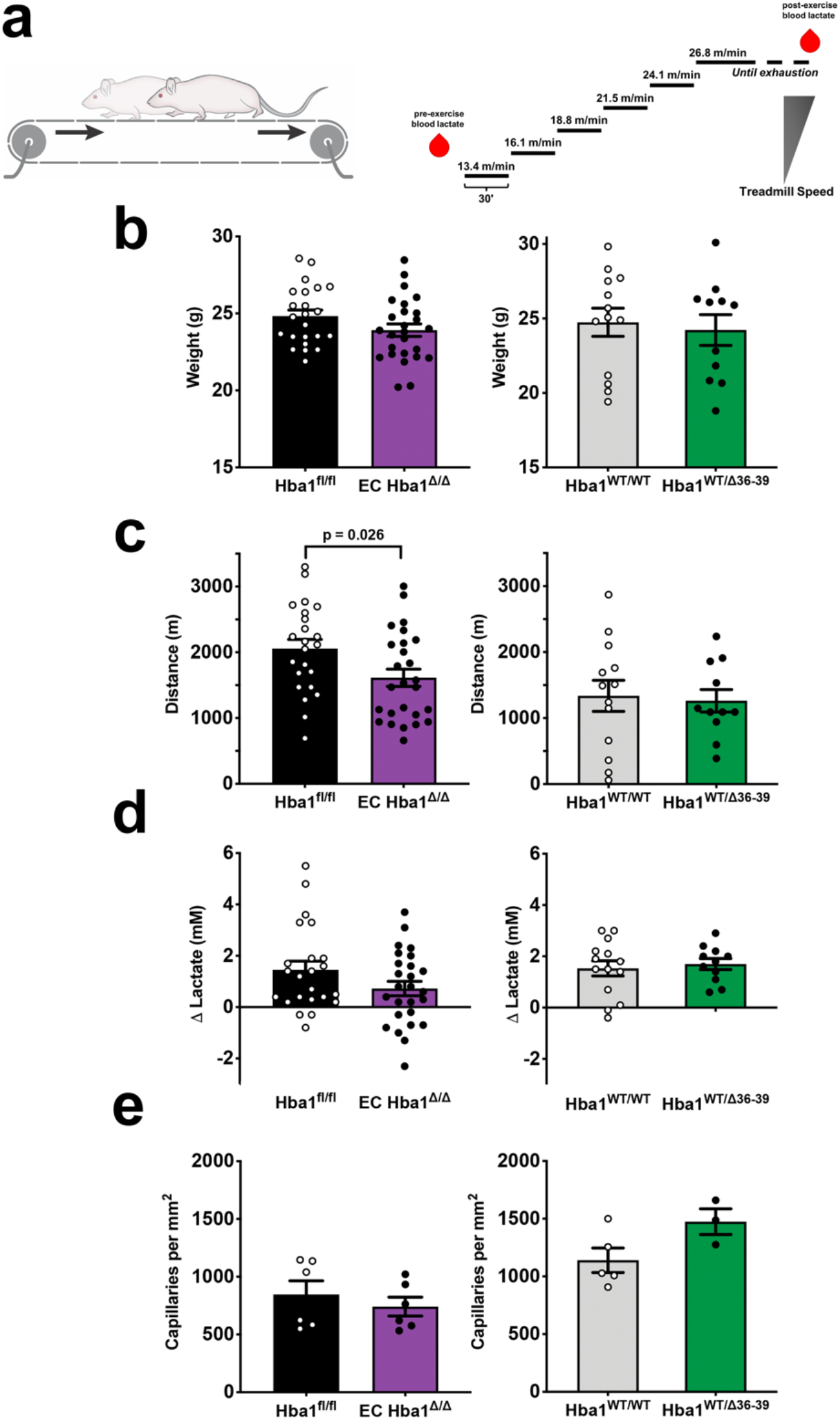
Exercise capacity is reduced with total EC alpha globin loss of function, but not with eNOS binding disruption. (**a**) Schematic of the exercise capacity protocol. After an initial blood sample for baseline lactate measurement, mice are encouraged to run on a treadmill until exhaustion. Speed of the treadmill is increased every 30 minutes until a maximum speed of 26.8 m/min (1 mile per hour) is achieved. After running failure, a second blood sample is taken for lactate buildup monitoring to ensure physical exhaustion. (**b**) Body weight measurements for *Hba1*^*fl/fl*^ vs. EC *Hba1*^*Δ/Δ*^ or the *Hba1*^*WT/WT*^ vs. *Hba1*^*WT/Δ^36-39^*^ littermate groups were not different. (**c**) Distance to exhaustion was decreased in EC *Hba1*^*Δ/Δ*^ mice compared to *Hba1*^*fl/fl*^ littermates; this difference was not observed when comparing the distance to exhaustion of the *Hba1*^*WT/WT*^ and *Hba1*^*WT/Δ^36-39^*^ littermate groups. (**d**) Blood lactate was increased in all groups after exercise, but no differences were observed in between control and experimental groups. (**e**) Soleus muscle capillary density was similar across littermate comparisons. For b-d, n = 23 for *Hba1*^*fl/fl*^; n = 26 for EC *Hba1*^*Δ/Δ*^; n = 13 for *Hba1*^*WT/WT*^; and n = 11 for *Hba1*^*WT/Δ^36-39^*^. For e, n = 6 for *Hba1*^*fl/fl*^; n = 6 for EC *Hba1*^*Δ/Δ*^; n = 5 for *Hba1*^*WT/WT*^; and n = 3 for *Hba1*^*WT/Δ^36-39^*^. Comparisons between littermate genotypes were via a t-test.

### Endothelial alpha globin controls vessel diameter through nitrite reduction

To determine whether endothelial alpha globin can act as a nitrite reductase and affect individual arterial responses to hypoxia, we measured intracellular nitrite and nitrate concentrations from isolated thoracodorsal arteries. Increasing doses of sodium dithionite (Na_2_S_2_O_4_) were incubated with the isolated arteries to recapitulate the hypoxia experienced by skeletal muscle arteries in exercise (**Supplemental Figure 10**). From control vessels, we found a significant decrease in nitrite concurrent with an increase in cellular nitrate (**Figure 5**). In the EC *Hba1*^*Δ/Δ*^ vessels, we observe nitrite and nitrate concentrations that match untreated control vessels and are significantly different from littermate *Hba1*^*fl/fl*^ mice. These data indicate that the isolated artery itself, independent of erythrocytes, is capable of consuming nitrite in response to hypoxia, and that the consumption of nitrite is correlated with presence of endothelial alpha globin.

The vasodilatory response of isolated arteries induced by hypoxia was measured using pressure myography (**Figure 6A**). In genotypes where endothelial alpha globin is intact and can participate in hypoxic nitrite reduction (WT, *Hba1*^*fl/fl*^ and both *Hba1*^*Δ36-39*^ genotypes), a robust dilation in response to Na_2_S_2_O_4_ is observed. However, in the EC *Hba1*^*Δ/Δ*^ group, vasodilation is blunted across increasing doses of Na2S2O4 (**Figure 6B**), an observation that remains consistent with global deletion of *Hba1* (**Supplemental Figure 11**). Other experiments with pharmacologic agents were performed with a single dose of 1 mM Na_2_S_2_O_4_, as it is in the middle of our range and produced a robust dilation (**Figure 6C**).

**FIGURE 5:**
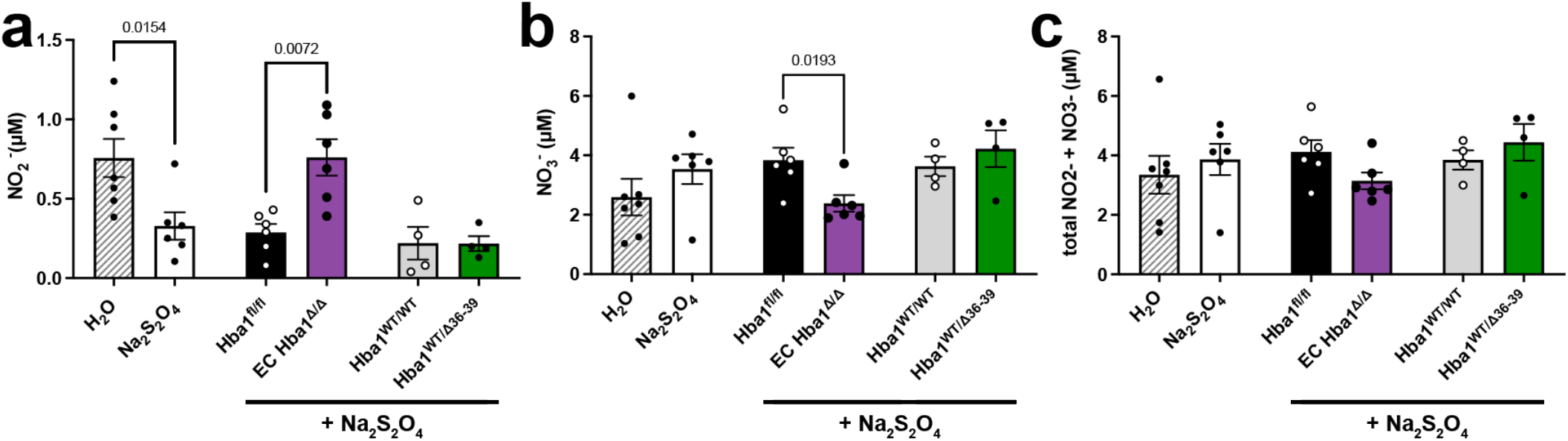
Nitrite consumption is increased by chemical hypoxia in isolated thoracodorsal vessels. Nitrite (a), nitrate (b), and total NO_x_ species (c) were measured after incubation of an isolated vessel with sodium dithionite (Na_2_S_2_O_4_). WT thoracodorsal vessels treated with Na_2_S_2_O_4_ show decreased intracellular nitrite compared to vessels treated with water (H_2_O) (leftmost group). All experimental genotypes were treated with Na_2_S_2_O_4_. EC *Hba1*^*Δ/Δ*^ showed higher levels of nitrite and lower nitrate levels compared to littermate controls with chemical hypoxia, while total NO_x_ was not different. For H_2_O treatment, n = 7; for WT vessels treated with Na_2_S_2_O_4_, n = 6; for *Hba1*^*fl/fl*^ n = 6; for EC Hba1^Δ/Δ^ n = 6; for *Hba1*^*WT/WT*^ n = 4; and n = 4 for *Hba1*^*WT/Δ^36-39^*^. Comparisons made between littermates via a t-test.

**FIGURE 6:**
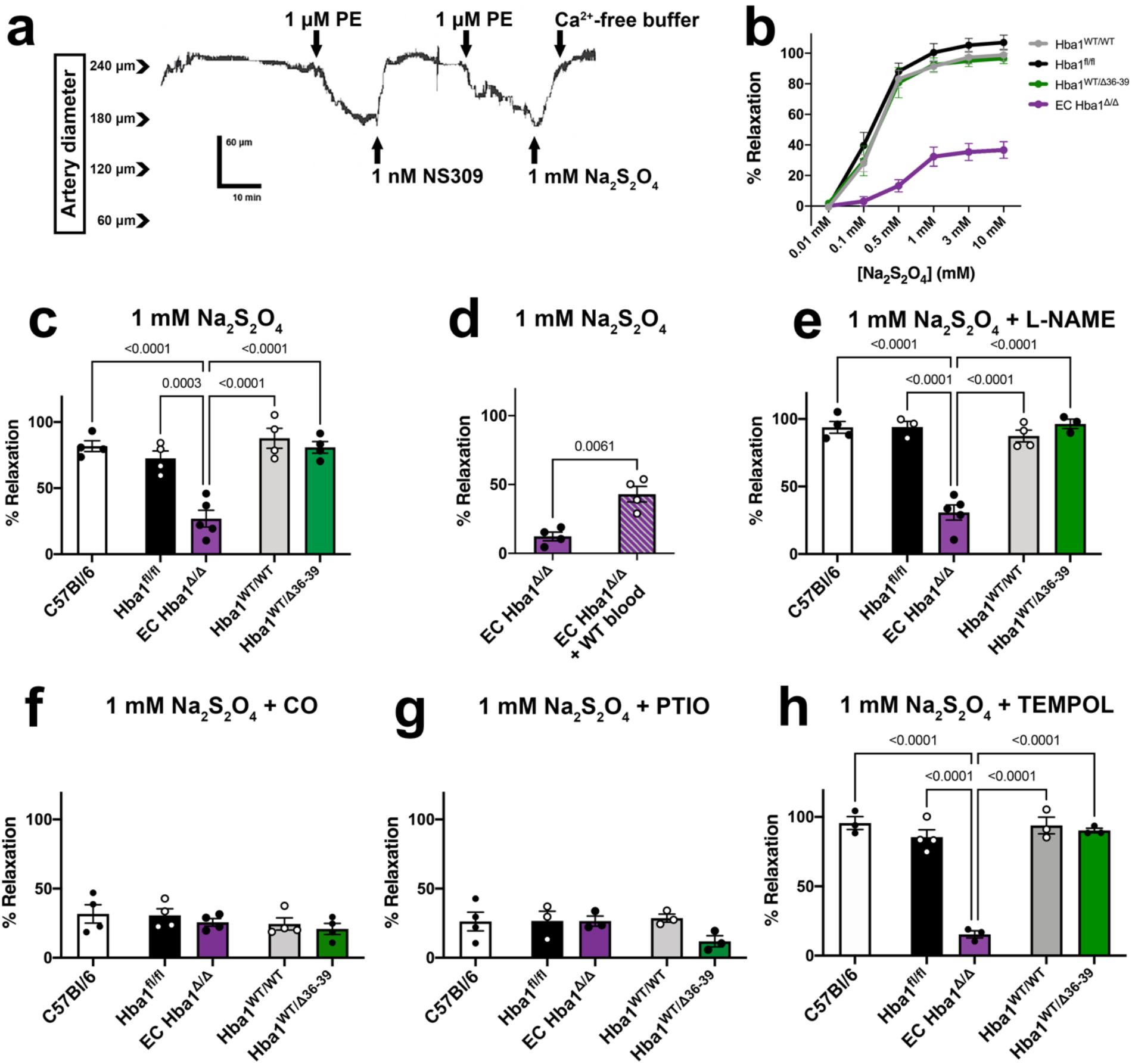
Hypoxic vasodilation is decreased without endothelial alpha globin. (**a**) Representative trace of hypoxic vasodilation experiment. Cannulated vessels are treated with an initial dose of phenylephrine (PE) to constrict, and endothelial dilation integrity confirmed by dilation to NS309. Another PE dose constricts, and dilation to sodium dithionite (Na_2_S_2_O_4_) is assessed. Finally, maximal dilation to Ca^2+^-free buffer establishes the percent of maximal dilation for each treatment. Vasodilation in response increasing doses of Na_2_S_2_O_4_ is blunted by endothelial deletion of alpha globin (purple), but not by decreasing its interaction with eNOS (green). (**c**) Hypoxia induced by a single dose of 1 mM Na_2_S_2_O_4_ shows blunted dilation only in the EC *Hba1*^*Δ/Δ*^ genotype. The hypoxic vasodilation is preserved in *Hba1*^*fl/fl*^, *Hba1*^*WT/WT*^, and *Hba1*^*WT/Δ^36-39^*^ groups. (**d**) Addition of blood to the lumen of the EC *Hba1*^*Δ/Δ*^ vessel partially restores dilation after Na_2_S_2_O_4_ treatment. (**e**) The hypoxic vasodilation response is not affected by pretreatment of the vessel with L-NAME, a NOS inhibitor. (**f**) This hypoxic vasodilation response of the isolated resistance artery is inhibited by carbon monoxide. (**g**) The dilation in response to Na_2_S_2_O_4_ is NO-mediated, as treatment with the NO scavenger PTIO (2-phenyl-4, 4, 5, 5,-tetramethylimidazoline-1-oxyl 3-oxide) prevents dilation in all groups. (**h**) Limiting other reactive oxygen species (including superoxide and hydrogen peroxide) with TEMPOL does not restore a dilatory response to the EC *Hba1*^*Δ/Δ*^ group in response to chemical hypoxia. All experiments were performed with at least n > 3 separate animals per genotype. All comparisons between groups were made with a t-test with multiple comparisons correction, except in d, where a t-test was used.

As a control, we reintroduced red blood cells into the lumen of EC *Hba1*^*Δ/Δ*^ arteries to determine if erythrocyte-dependent nitrite reduction was sufficient to restore dilatory response to Na_2_S_2_O_4_-induced hypoxia. A statistically significant, yet partial, rescue of vasodilation was observed from luminal erythrocytes in Na_2_S_2_O_4_ treatment (**Figure 6D**).

To block NO production from NOS, isolated arteries were incubated with the NOS inhibitor L-nitroarginine methyl ester (L-NAME) prior to Na_2_S_2_O_4_-induced hypoxia (**Figure 6E**). NOS inhibition did not affect the robust dilation for the *Hba1*^*fl/fl*^, *Hba1*^*WT/WT*^, or *Hba1*^*WT/Δ36-39*^ mice. Dilation of the arteries of EC *Hba1*^*Δ/Δ*^ mice remained low in this context; thus, the hypoxic vasodilation response appears NOS-independent but relies on the presence of the full alpha globin protein.

Nitrite reduction is dependent on heme redox chemistry. To prevent nitrite reduction by endothelial alpha globin heme iron, we pre-incubated the isolated vessel with carbon monoxide (CO), which tightly binds the heme group and prevents heme-based catalysis and nitrite reduction (**Figure 6F**). Treatment with CO fully blocked hypoxic vasodilation responses of all groups to levels equivalent to EC *Hba1*^*Δ/Δ*^ mice; thus, the hypoxic vasodilation is dependent on redox chemistry that can be inhibited by CO. Further, incubation of the vessel with the NO scavenger PTIO (2-phenyl-4, 4, 5, 5,-tetramethylimidazoline-1-oxyl 3-oxide) blunts dilation in all groups, demonstrating that the hypoxic vasodilation is dependent on NO signaling (**Figure 6G**).

Lastly, other vasoactive molecules produced by endothelium might contribute to the hypoxic vasodilation response. Hydrogen peroxide or other reactive oxygen species could cause dilation in hypoxia; treatment of the isolated vessel with the peroxide scavenger TEMPOL did not affect baseline dilation of any genotype (**Figure 6H**), excluding hydrogen peroxide as the vasodilatory signal in this context.

### Blood pressure change to environmental hypoxia is blunted in EC Hba1^Δ/Δ^ mice

Hypoxic vasodilation can also influence arterial blood pressure. Using radiotelemetry, we monitored blood pressure of the *Hba1*^*fl/fl*^, EC *Hba1*^*Δ/Δ*^, *Hba1*^*WT/WT*^, and *Hba1*^*WT/Δ36-39*^ mice. When exposed to room air (normoxia), the EC *Hba1*^*Δ/Δ*^ blood pressure was not significantly altered (**Figure 7A**). However, in environmental hypoxia (10% O_2_), the *Hba1*^*fl/fl*^, *Hba1*^*WT/WT*^, and *Hba1*^*WT/Δ36-39*^ mice exhibited decreased systolic blood pressure compared to normoxic baseline, indicating a global vasodilatory response (as observed before^34^) (**Figure 7B**). However, the blood pressure in EC *Hba1*^*Δ/Δ*^ mice remained relatively unchanged (112 mmHg in normoxia to 108 mmHg in hypoxia), consistent with the observed lack of hypoxic vasodilation response in isolated arteries. The difference between normoxic and hypoxic systolic blood pressures are quantified in **Figure 7C**. Compared to littermate controls, the EC *Hba1*^*Δ/Δ*^ genotype show significantly less difference in blood pressure between normoxia and hypoxia.

**FIGURE 7:**
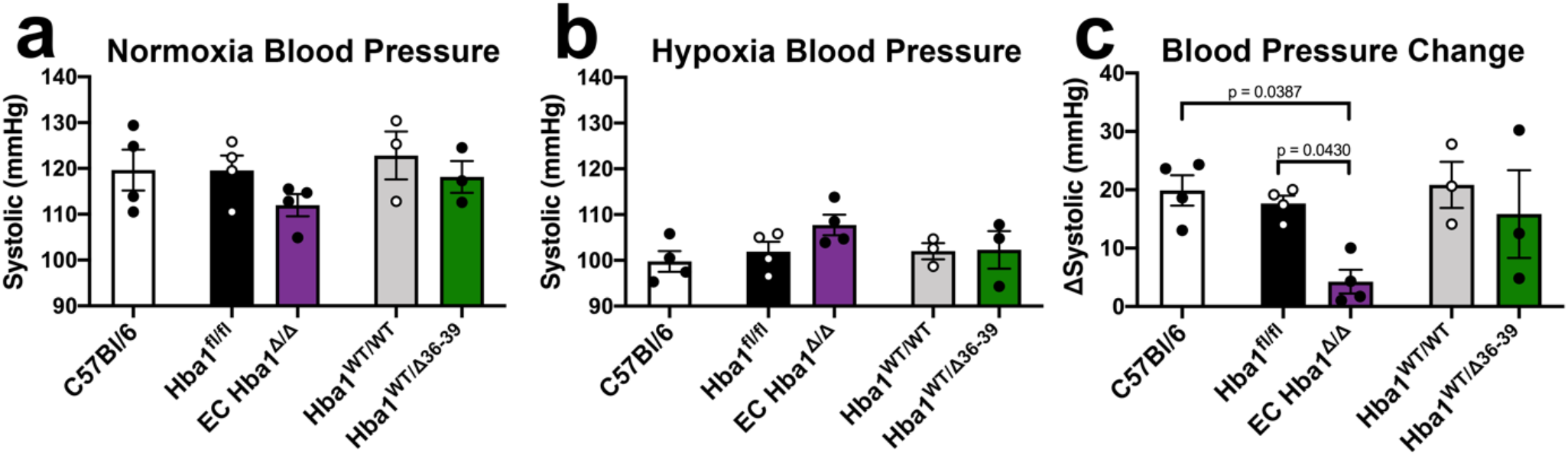
Endothelial alpha globin enhances blood pressure change to global hypoxia. (**a**) In normoxia (room air) the systolic blood pressure of the EC *Hba1*^*Δ/Δ*^ genotype is slightly decreased. (**b**) When exposed to hypoxia, the EC *Hba1*^*Δ/Δ*^ group has slightly elevated systolic blood pressure compared to other groups. (**c**) EC *Hba1*^*Δ/Δ*^ mice have the smallest change in systolic blood pressure difference between normoxia and hypoxia. This effect is statistically different from littermate controls and WT mice. For C57Bl/6 mice, n = 4; for *Hba1*^*fl/fl*^ n = 3; for EC *Hba1*^*Δ/Δ*^ n = 4; for *Hba1*^*WT/WT*^ n = 3; and n = 3 for *Hba1*^*WT/Δ^36-39^*^. Comparisons were made across groups using a one-way ANOVA.

## Discussion

Hypoxic vasodilation through NO generation from hemoproteins has been described previously^35^ and there is strong evidence of its significant role in the regulation of circulatory physiology in general. Erythrocytic deoxygenated hemoglobin can act as a nitrite reductase to provide NO as a vasodilation signal to vessels. However, the circulatory location where this signaling occurs (i.e. resistance arteries vs. post-capillary venules^36^) and the efficiency of erythrocyte NO in reaching vascular smooth muscle^5^ remain open questions. We hypothesized that endothelial alpha globin is uniquely situated to provide NO to vascular smooth muscle in hypoxic contexts. Our study identifies a novel nitrite reductase function for endothelial alpha globin that may contribute to hypoxic vasodilation.

A role for alpha globin as a nitrite reductase presents an interesting juxtaposition with prior reports of alpha globin inhibiting NO signaling through interaction with eNOS and NO scavenging^9, 10, 11, 37^. However, the two roles for alpha globin are distinct in the metabolic contexts in which they dominate. Alpha globin inhibiting eNOS-derived NO signaling necessarily requires O_2_, as both eNOS enzyme activity and NO scavenging and subsequent transformation to NO_3_^−^ by alpha globin require O_2_. On the other hand, alpha globin (as with erythrocyte hemoglobin) must be deoxygenated to act as a nitrite reductase^36^. Thus, the opposed roles for alpha globin in NO signaling are separated by local metabolism: when O_2_ is prevalent, alpha globin may predominate as an NO signaling inhibitor. However, when tissues are metabolically active and O_2_ is scarce, endothelial alpha globin can act as a hypoxia sensor and signal enactor to increase perfusion and match O_2_ supply to metabolic demand.

To determine whether alpha globin could control vasodilation through nitrite reduction separate from its role in eNOS-derived NO scavenging, we used two novel loss-of-function mouse models to demonstrate that endothelial alpha globin participates in hypoxic vasodilation through an autonomous nitrite reductase mechanism. First, an endothelial-specific deletion of alpha globin (EC *Hba1*^*Δ/Δ*^) demonstrates the effects of loss of eNOS-inhibition and nitrite reductase (through the heme moiety) functions of endothelial alpha globin. Second, the global mutation of alpha globin through deletion of four residues (*Hba1*^*Δ36-39*^) prevents the association of alpha globin and eNOS, thereby decreasing NO scavenging^9, 10^, while preserving nitrite reductase activity. With the combination of these two models, we are able to specifically assay a nitrite reductase role for endothelial alpha globin.

First, the distance to exhaustion assayed by the treadmill running test was decreased in the EC *Hba1*^*Δ/Δ*^ genotype compared to littermate *Hba1*^*fl/fl*^ controls (**Figure 4**). This difference is in contrast to the *Hba1*^*WT/WT*^ and *Hba1*^*WT/Δ36-39*^ comparison, where the animals were exhausted at a similar distance between littermates. Additionally, in the global *Hba1*^*-/-*^ genotype, distance to exhaustion was also decreased, although this is confounded by the anemia in that genotype (**Supplemental Figure 9** and **Supplemental Figure 4**). Running distance appears different between *Hba1*^*fl/fl*^ genotypes and *Hba1*^*WT/Δ36-39*^ genotypes, although this discrepancy could be due to strain or background differences.

Second, vascular cell nitrite utilization appears significantly decreased in EC *Hba1*^*Δ/Δ*^ mice (**Figure 5**). Incubation of isolated thoracodorsal arteries with Na_2_S_2_O_4_ resulted in WT vessels having significantly decreased nitrite. Vessels isolated from EC *Hba1*^*Δ/Δ*^ mice had nitrite levels similar to untreated vessels, even after 5 minutes of Na_2_S_2_O_4_ treatment. There was a significant increased level of cellular nitrite in EC *Hba1*^*Δ/Δ*^ mice compared to littermates, while the *Hba1*^*Δ36-39*^ genotypes were similar to each other. Additionally, EC *Hba1*^*Δ/Δ*^ mice had decreased nitrate levels with unchanged total NO_x_ species. Overall, utilization of cellular nitrite in the thoracodorsal artery is significantly affected by the presence of endothelial alpha globin.

Finally, a dilatory response to hypoxia is absent in EC *Hba1*^*Δ/Δ*^ vessels, where endothelial alpha globin is absent. Loss of the entire alpha globin protein in either EC *Hba1*^*Δ/Δ*^ or global *Hba1*^*-/-*^ resistance vessels largely prevented dilation to hypoxic buffer conditions (**Figure 6B and Supplemental Figure 11**). However, targeted deletion of the eNOS-interacting motif did not change the vasodilation response to hypoxia. We determined that the vasodilatory response was NOS- and hydrogen peroxide-independent, while requiring NO. Furthermore, the dilation to Na_2_S_2_O_4_-induced hypoxia is dependent on a CO-inhibitable process, consistent with heme-based nitrite reduction. Integrating the dilatory response of individual arteries with whole animal physiology, the normal decrease in blood pressure associated with arterial hypoxia was abrogated in mice lacking endothelial alpha globin expression. The only differences observed in these experiments occurred when endothelial alpha globin was deleted and thus unable to participate in nitrite reduction.

We acknowledge a few limitations in these studies. First, the *Hba1*^*Δ36-39*^ model is a complicated model due to homozygous lethality; thus, heterozygous adults were necessarily used in the experiments on adult animals. Additionally, this mutation is constitutive and global, precluding observations on endothelial alpha globin specifically. However, the use of littermate controls in all studies allows for phenotypic comparisons of WT and heterozygous disruption of alpha globin/eNOS interaction. Second, many of our experiments were performed in a whole animal physiology context where erythrocytes can contribute to hypoxic vasodilation. Although our EC *Hba1*^*Δ/Δ*^ mutation is targeted to endothelium using the *Cdh5-Cre*^*ERT2*^, we did not specifically measure the contribution of erythrocytes to NO signaling in exercise hyperemia or hypoxic blood pressure compensation. The use of isolated skeletal muscle resistance arteries from each of our genetic contexts demonstrated that endothelial alpha globin can induce vasodilation, but the proportions of endothelial alpha globin or erythrocytic hemoglobin signaling were not directly measured.

Overall, our experiments demonstrate that endothelial alpha globin can participate in hypoxic vasodilation through nitrite reduction. Alpha globin has an optimal expression localization that is critical for its function in controlling hypoxic vasodilation. First, endothelial alpha globin expression is normally restricted to resistance vasculature, specifically arterioles^11^. Thus, as a nitrite reductase, alpha globin can provide NO signaling where it is maximally effective to enact dilation and changes in tissue perfusion. Dilation of the pre-capillary arterial beds increases tissue perfusion without the need for signals to propagate up or down the vascular tree. Second, the subcellular localization of alpha globin to the myoendothelial junction (MEJ)^11, 37^ could provide additional benefits in spatial optimization of NO generation. The MEJ is a domain of the endothelium that is in direct contact with underlying vascular smooth muscle^38^, so generation of NO by MEJ-restricted alpha globin minimizes the diffusion distance necessary to enact vasodilation through smooth muscle relaxation.

This novel role for endothelial alpha globin places alpha globin at a unique fulcrum of NO signaling in the resistance vasculature. As both an inhibitor of eNOS-produced NO and a source of NO generation in hypoxia, alpha globin uniquely determines resistance vascular function in the endothelium.

## Acknowledgments

This work was supported by NIH HL088554 (BEI and MW), GM100776 (TPB), HL098032 (DBKS), R35 GM131829 (LC), F31HL1440032 (TCSK). This research was supported in part by the Intramural Research Program of the NIH, National Institute of Allergy and Infectious Diseases (SB and HA).

Authors declare no financial interest.

## Supplemental Figures

**SUPPLEMENTAL FIGURE 1:**
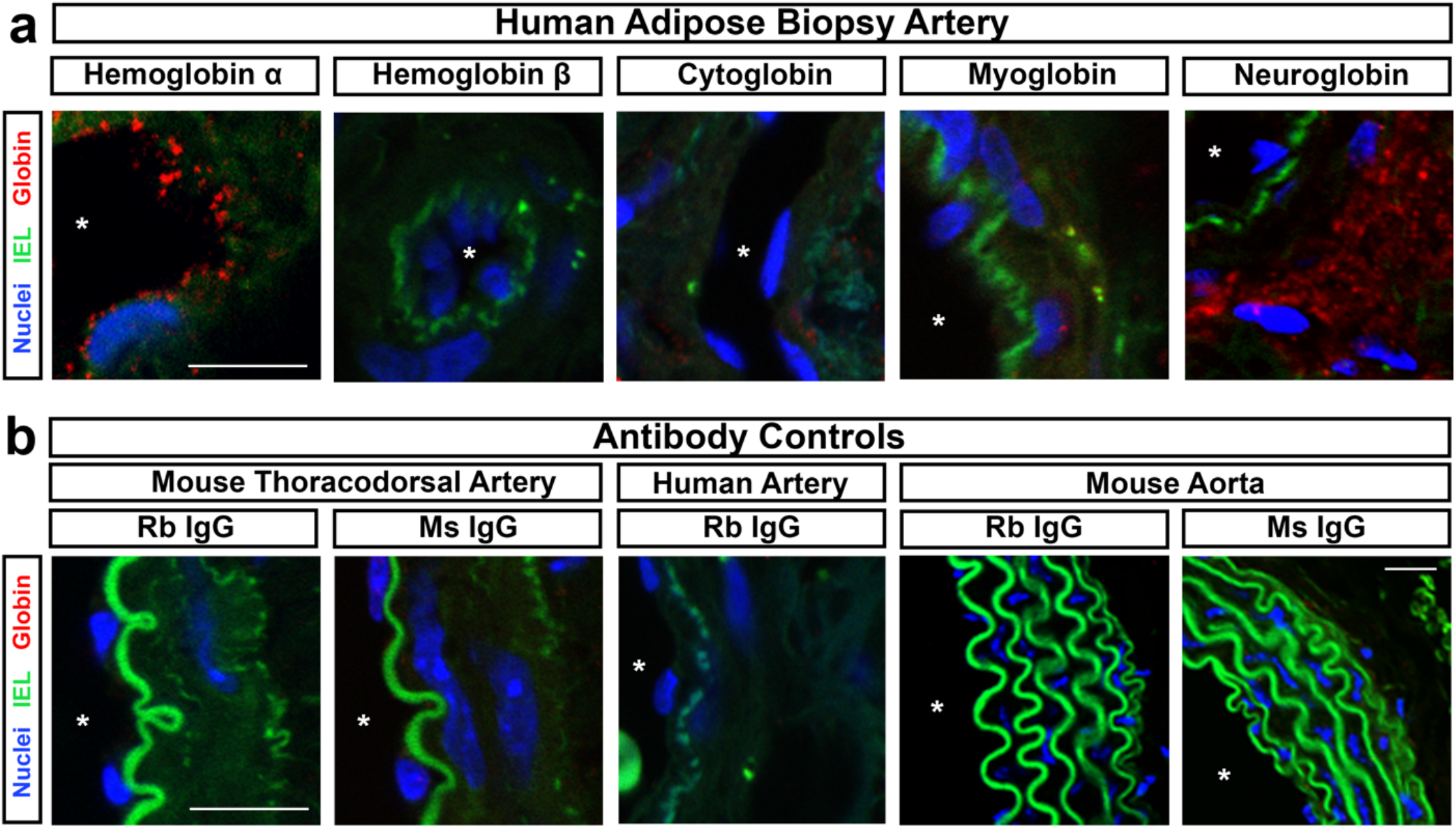
Alpha globin is expressed in the endothelium of arterioles from human adipose biopsy. (A) Expression of alpha globin in the human endothelium is observed from adipose biopsy samples. No other globin is observed in the endothelium; neuroglobin signal is observed in the adventitial layers. (B) antibody controls for the positive signals observed in the tissues from Main Text Figure 1 and this figure are presented to demonstrate specificity of staining. In all images, an asterisk denotes the lumen of the vessel. Scale bars in a are 10 μm, and in b are 25 μm.

**SUPPLEMENTAL FIGURE 2:**
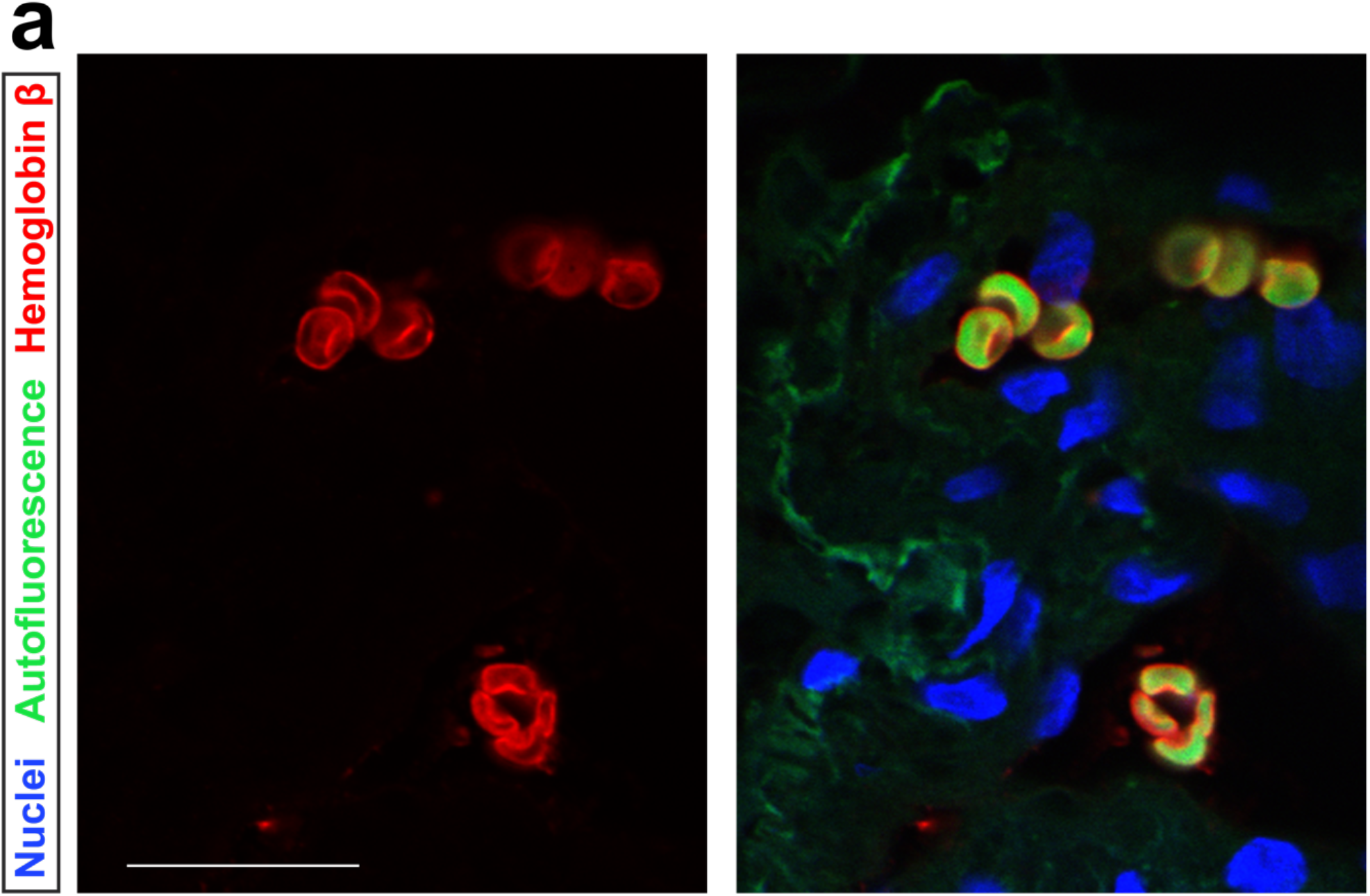
Staining for hemoglobin β is seen in erythrocytes, as an immunohistochemistry positive control. (**a**) Positive signal (red) for hemoglobin β is observed in erythrocytes captured in a section of human adipose biopsy. Erythrocytes are strongly autofluorescent (green), and nuclei are denoted with DAPI staining (blue). Scale bar is 30 μm.

**SUPPLEMENTAL FIGURE 3:**
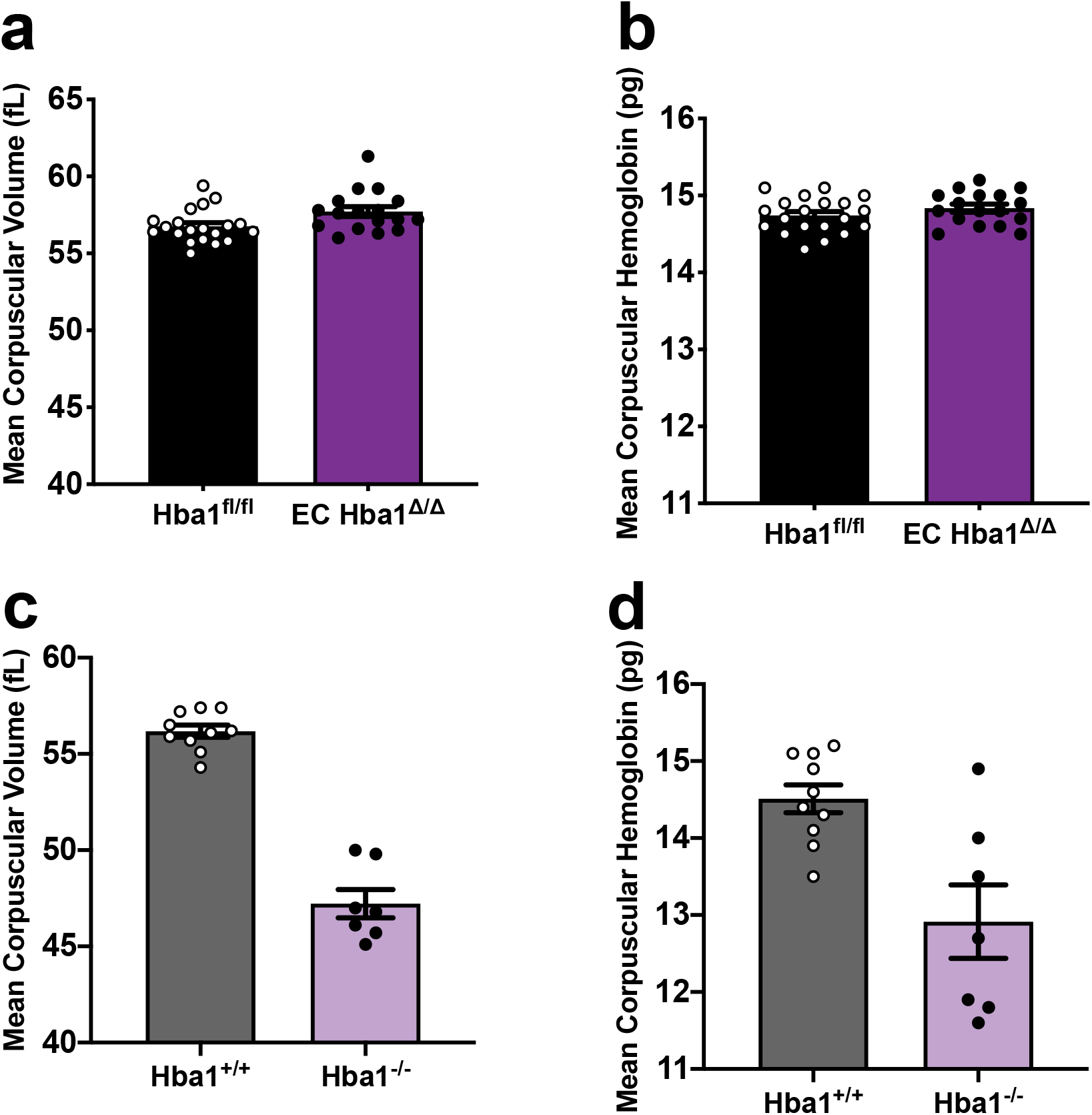
Other hematologic measurements for the EC Hba1^Δ/Δ^ model. (**a**) Mean corpuscular volume (MCV) or mean corpuscular hemoglobin (MCH) (**b**) are not changed with endothelial-specific (*Cdh5-Cre*^*ERT2*^) recombination of the *Hba1*^*fl/fl*^ allele. In the neomycin-insertion global deletion (*Hba1*^*-/-*^), both MCV (**c**) and MCH (**d**) are decreased compared to WT controls. For *Hba1*^*fl/fl*^ n = 20; for EC *Hba1*^*Δ/Δ*^ n = 17; for *Hba1*^*+/+*^ n = 10; and n = 7 for *Hba1*^*-/-*^.

**SUPPLEMENTAL FIGURE 4:**
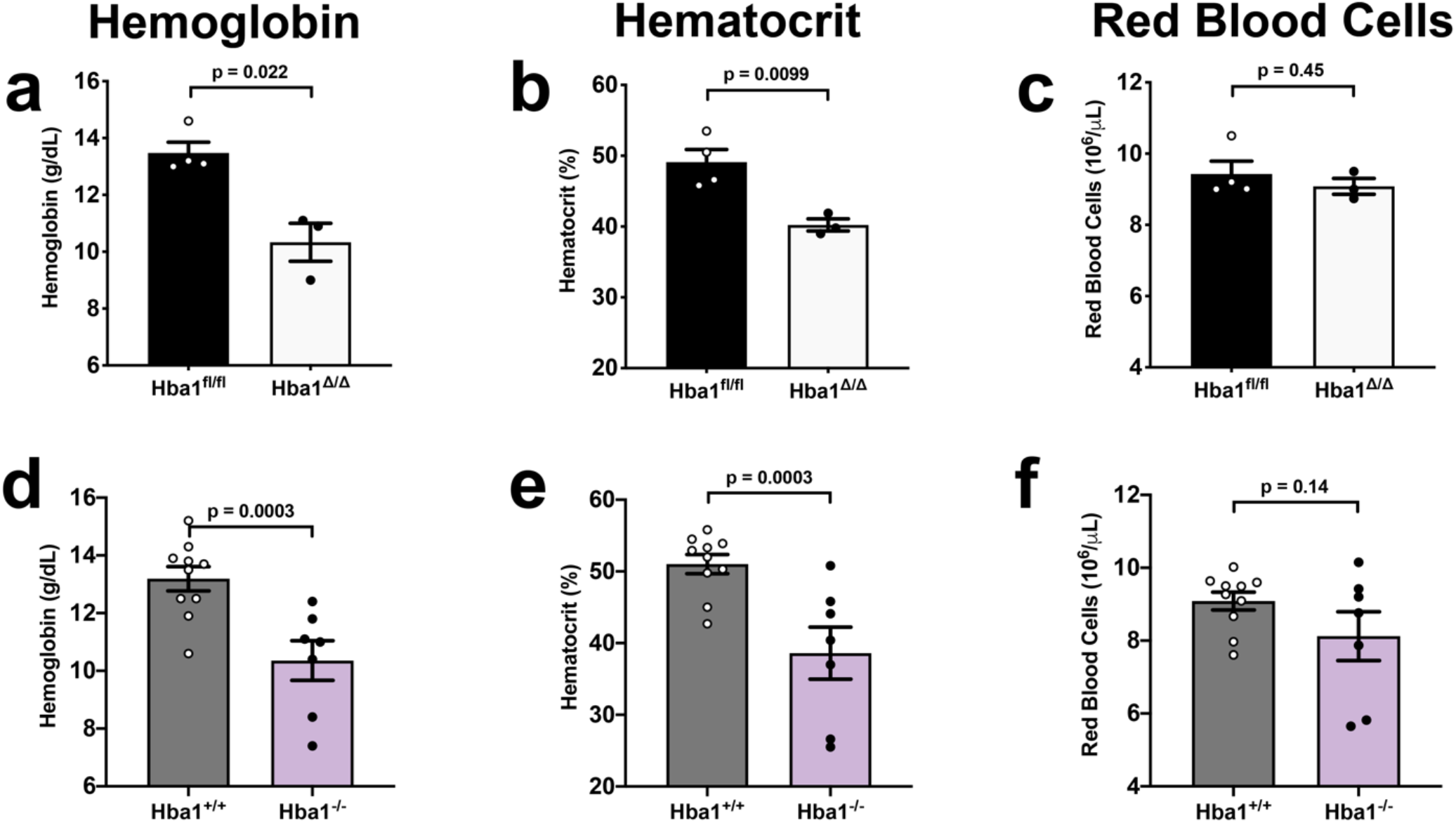
Blood parameters of mice with global deletion of Hba1 show anemic phenotype. Hemoglobin (**a, d**), hematocrit (**b, e**), and total RBC (**c, f**) show the same pattern in a global Hba1 KO (*Hba1*^*fl/fl*^; *Sox2-Cre*, denoted *Hba1*^*Δ/Δ*^) and a model of alpha thalassemia (*Hba1*^*-/-*^). For *Hba1*^*fl/fl*^ n = 4; for *Hba1*^*Δ/Δ*^ n = 3; for *Hba1*^*+/+*^ n = 10; and n = 7 for *Hba1*^*-/-*^. Comparisons were made with a t-test.

**SUPPLMENTAL FIGURE 5:**
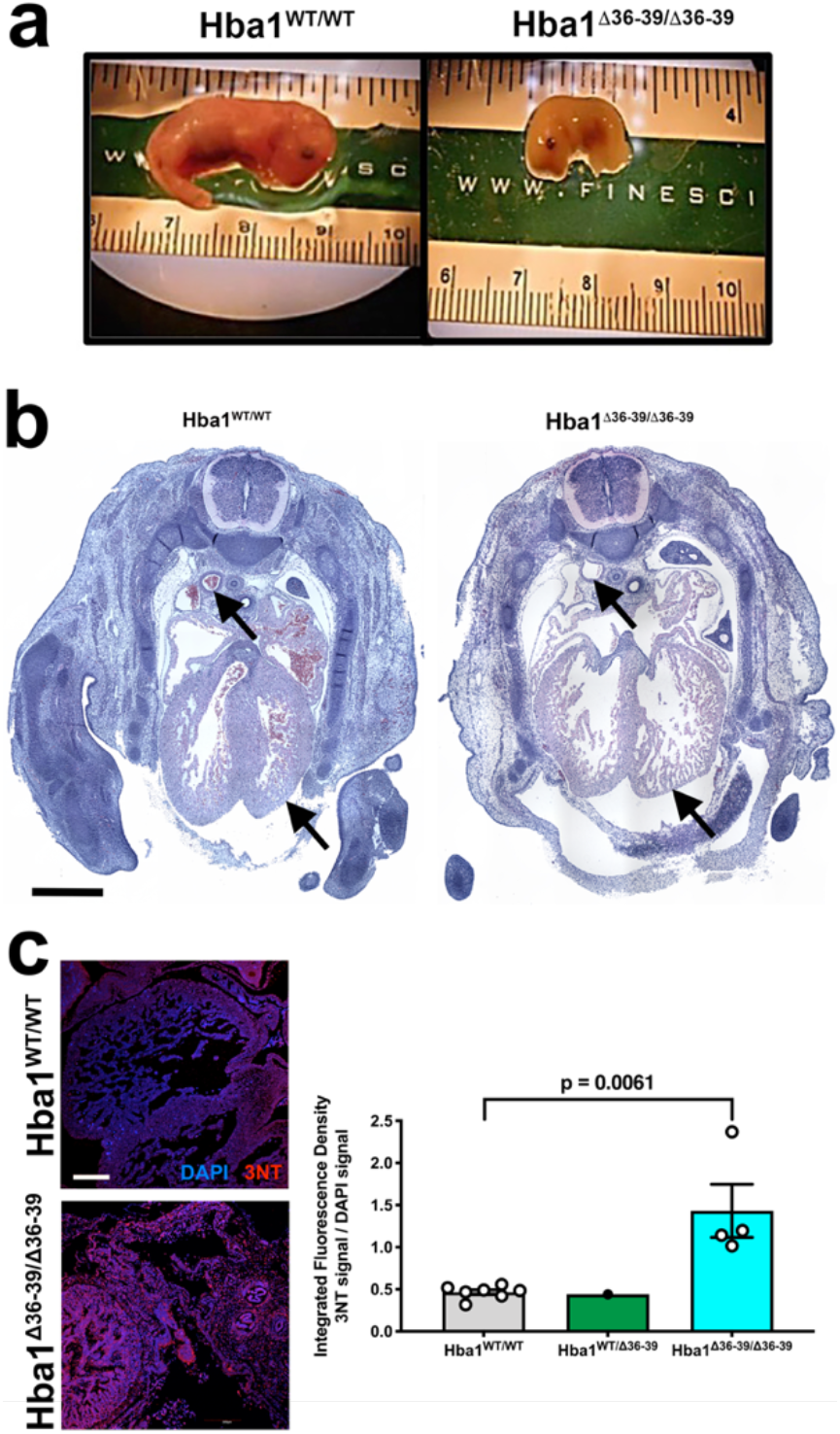
Homozygous mutation of Hba1^Δ36-39^ is embryonic lethal, with pathologic nitrotyrosine formation. Embryonic lethality is observed with homozygous Hba1^Δ36-39^ mutation. (**a**) Embryos at approximately E14.5 are already deceased with homozygous mutation, showing stunted growth. (**b**) Gross histology shows cardiac non-compaction, lack of blood, and dilated major vessels (arrows). Scale bare is 5 mm. (**c**) Immunostaining for 3-nitrotyrosine (a pathological marker of excess reactive nitrogen species), shows a large increase in signal in homozygous Hba1^Δ36-39^ mutation compared to WT or heterozygous embryos. Scale bar is 20 mm. For nitrotyrosine quantification in c, n = 7 for *Hba1*^*WT/WT*^, n = 1 for *Hba1*^*WT/Δ36-39*^, and n = 4 for *Hba1*^*Δ36-39/Δ36-39*^. Statistical comparison was made between *Hba1*^*WT/WT*^ and *Hba1*^*Δ36-39/Δ36-39*^ with a t-test.

**SUPPLEMENTAL FIGURE 6:**
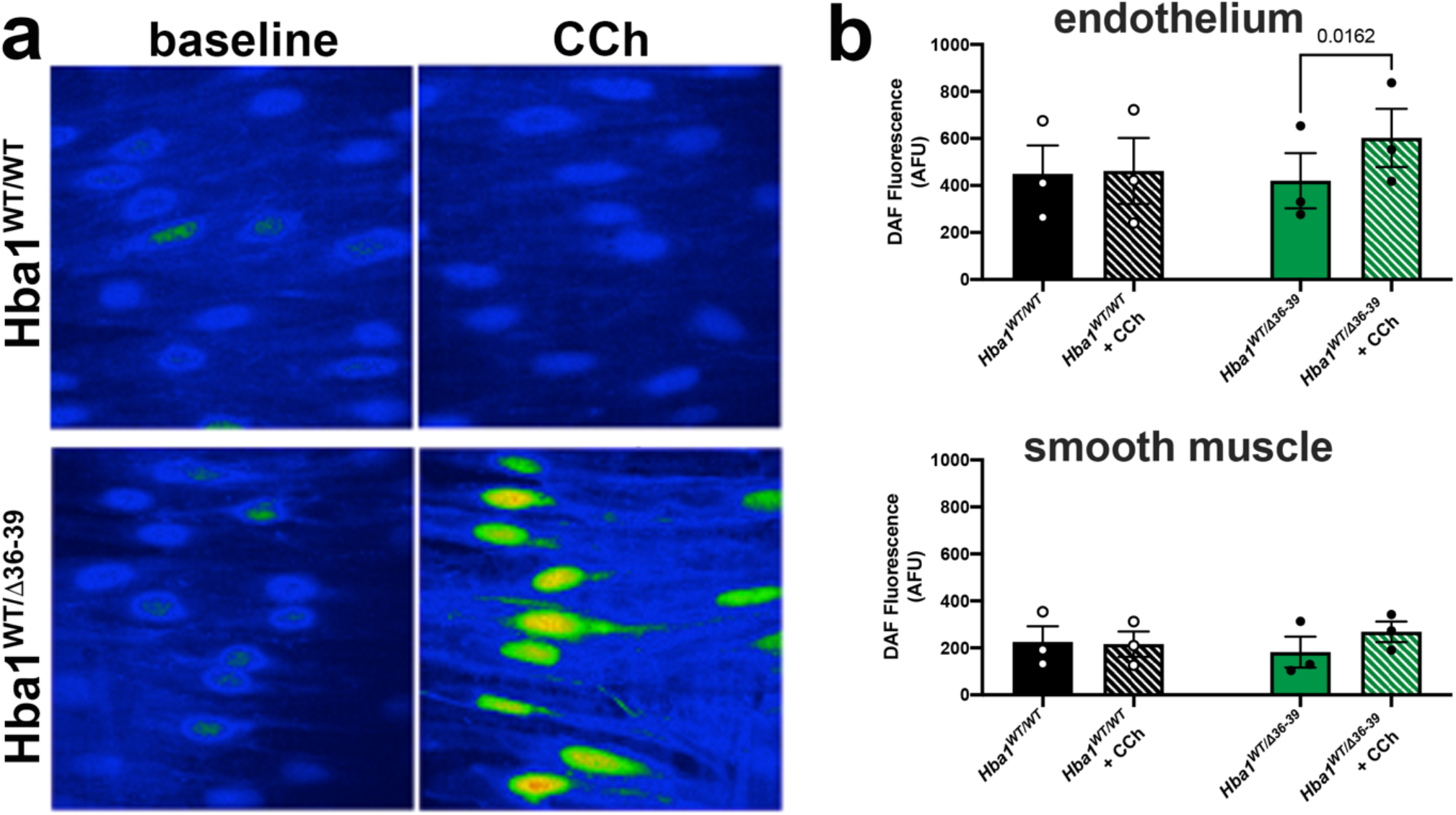
Increased nitric oxide generation in mice lacking an eNOS binding site in alpha globin. (**a**) *En face* imaging of third order mesenteric arterioles loaded with DAF were subject to 10 μM CCh stimulation and (**b**) the data quantified in the optical slices of endothelium or smooth muscle. N=3 mice for all measurements. Comparisons were made between littermate groups using a paired t-test. P-values are indicated where p<0.05.

**SUPPLEMENTAL FIGURE 7:**
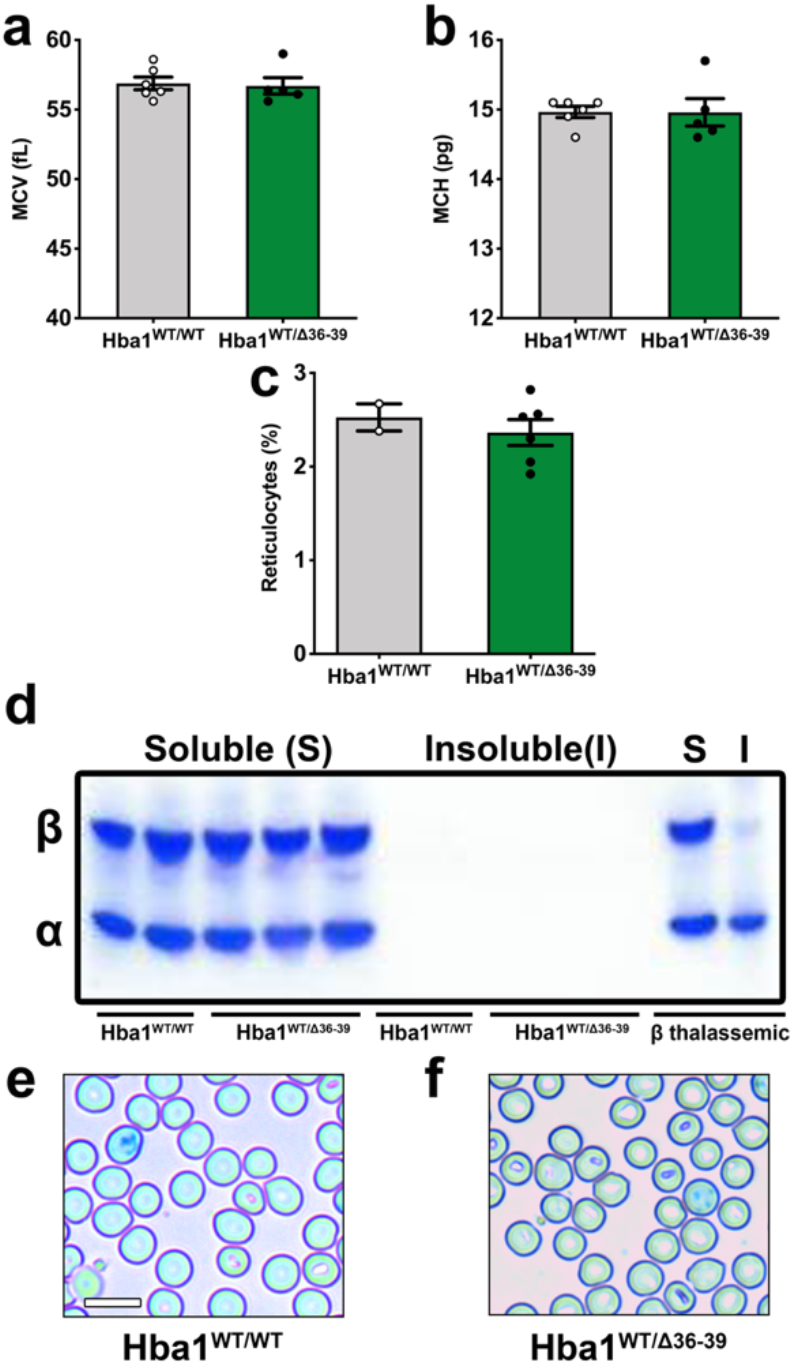
Hba1^WT/Δ36-39^ adults display unchanged red cell and reticulocyte characteristics, with no observed insoluble hemoglobin. (**a-c**) Blood cell parameters, including mean corpuscular volume (**a**), mean corpuscular hemoglobin (**b**), and number of reticulocytes (**c**), were not different between WT and heterozygous *Hba1*^*Δ36-39*^ mutants. (**d**) Blood samples from WT and *Hba1*^*Δ36-39*^ mutants do not show insoluble hemoglobin. A band for beta globin is at the top of the gel, and alpha globin is the lower band. Insoluble alpha globin is observed in a model of beta thalassemia (far right), as a control sample. (**e-f**) Cresyl blue staining of a blood smear demonstrates that heterozygous *Hba1*^*WT/Δ36-39*^ mutants do not have appreciably increased reticulocytes or precipitated hemoglobin compared to *Hba1*^*WT/WT*^ controls. Scale bar is 5 μm. For **a** and **b**, n = 6 for *Hba1*^*WT/WT*^, n = 5 for *Hba1*^*WT/Δ36-39*^, and for **c**, n = 2 for *Hba1*^*WT/WT*^, n = 6 for *Hba1*^*WT/Δ36-39*^. Comparisons between means were made with a t-test.

**SUPPLEMENTAL FIGURE 8:**
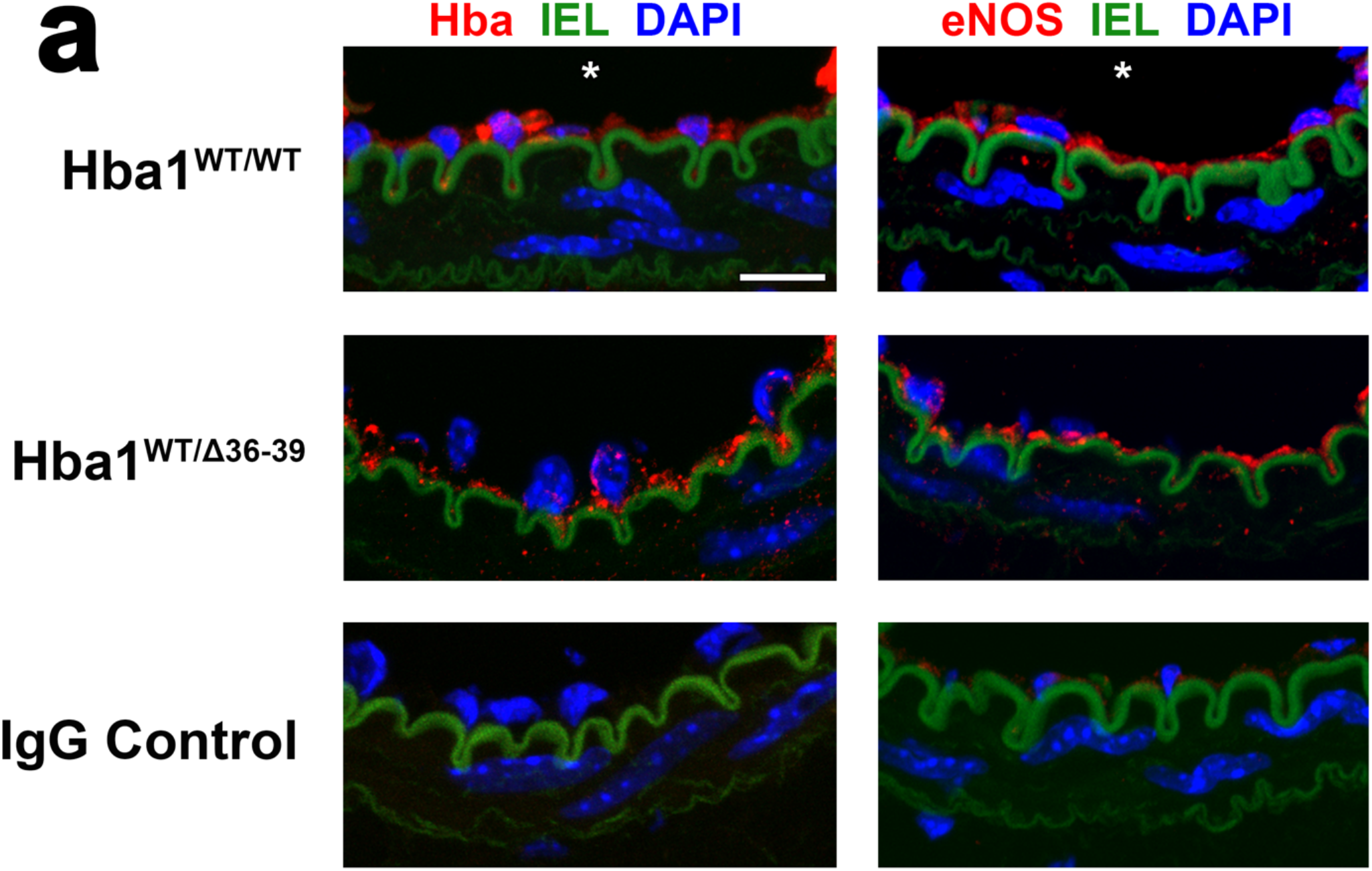
Alpha globin and eNOS are expressed to a similar level in Hba1^WT/WT^ and Hba1^WT/Δ36-39^ adult tissues. (a) Immunostaining controls for PLA. Alpha globin (left) and eNOS (right) are expressed and detectable to similar levels between *Hba1*^*WT/WT*^ and *Hba1*^*WT/Δ36-39*^ mutants in the thoracodorsal artery. Scale bar is 20 mm. The bottom row is stained with non-reactive IgG from the primary antibody host animal and shows no signal for either antigen.

**SUPPLEMENTAL FIGURE 9:**
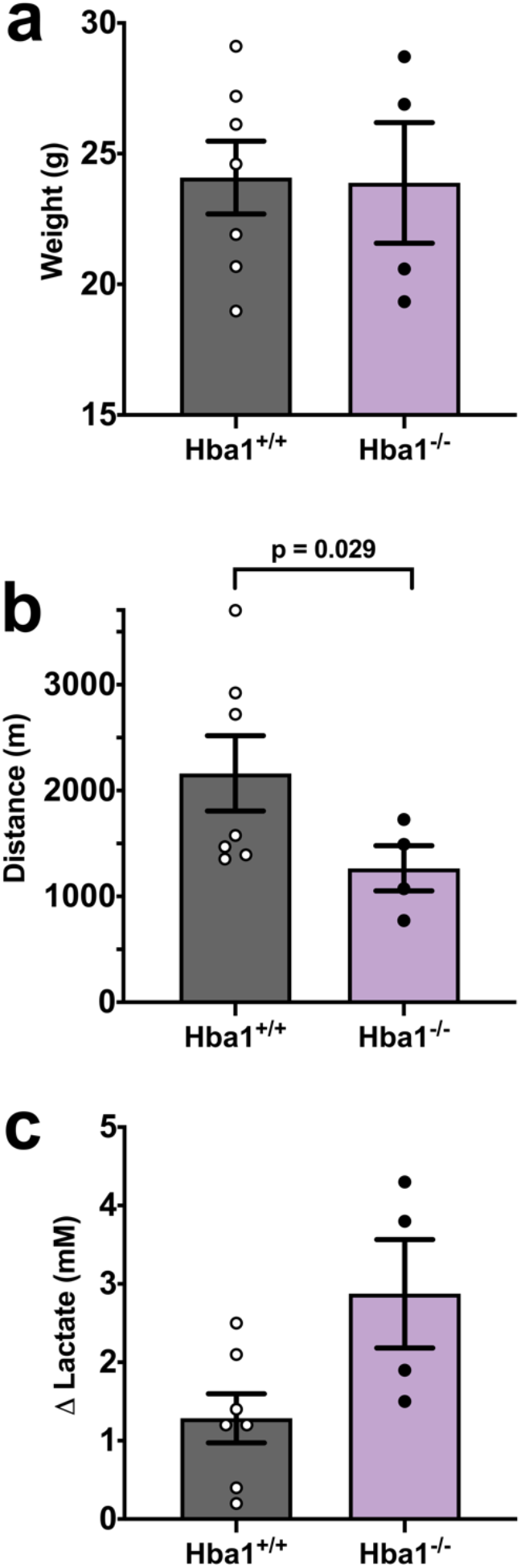
Global deletion of Hba1 reduces exercise capacity. Constitutive, global knockout of *Hba1* shows similar mouse weight (**a**) but decreased exercise capacity (**b**) with increases in blood lactate (**c**). The significant defect in running distance could be related to the anemia observed in these mice (Supplementary Figure 4). For *Hba1*^*+/+*^, n = 7 and for *Hba1*^*-/-*^, n = 4. Comparisons between groups were made with a t-test.

**SUPPLEMENTAL FIGURE 10:**
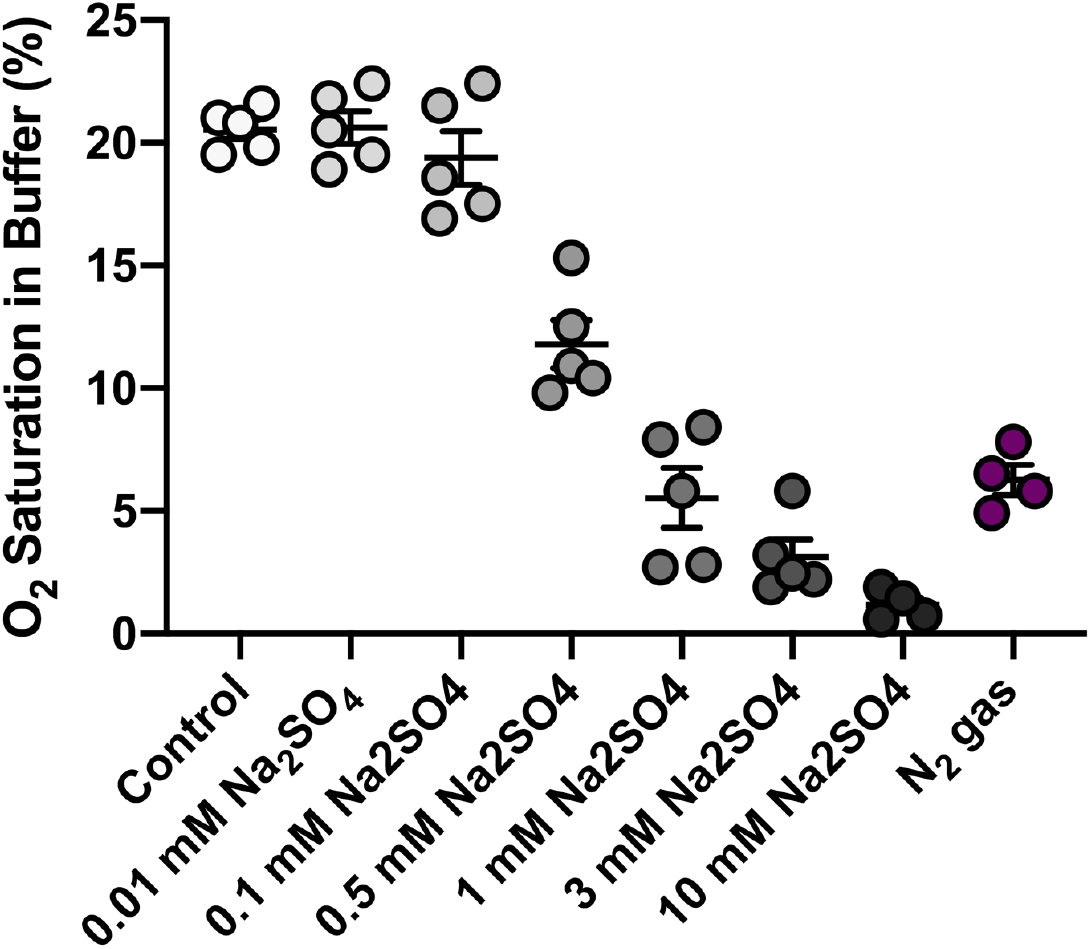
O_2_ content of buffer treatments with sodium dithionite used in the experiments. Increasing doses of Na_2_S_2_O_4_ decrease O_2_ saturation in the KREBS-based buffer used for vasoreactivity experiments. 1 mM Na_2_S_2_O_4_, used as the single dose in our studies, produces an aqueous O_2_ saturation similar to bubbling solution with pure nitrogen gas. For each condition, O_2_ saturation was measured in n = 5 samples.

**SUPPLEMENTAL FIGURE 11:**
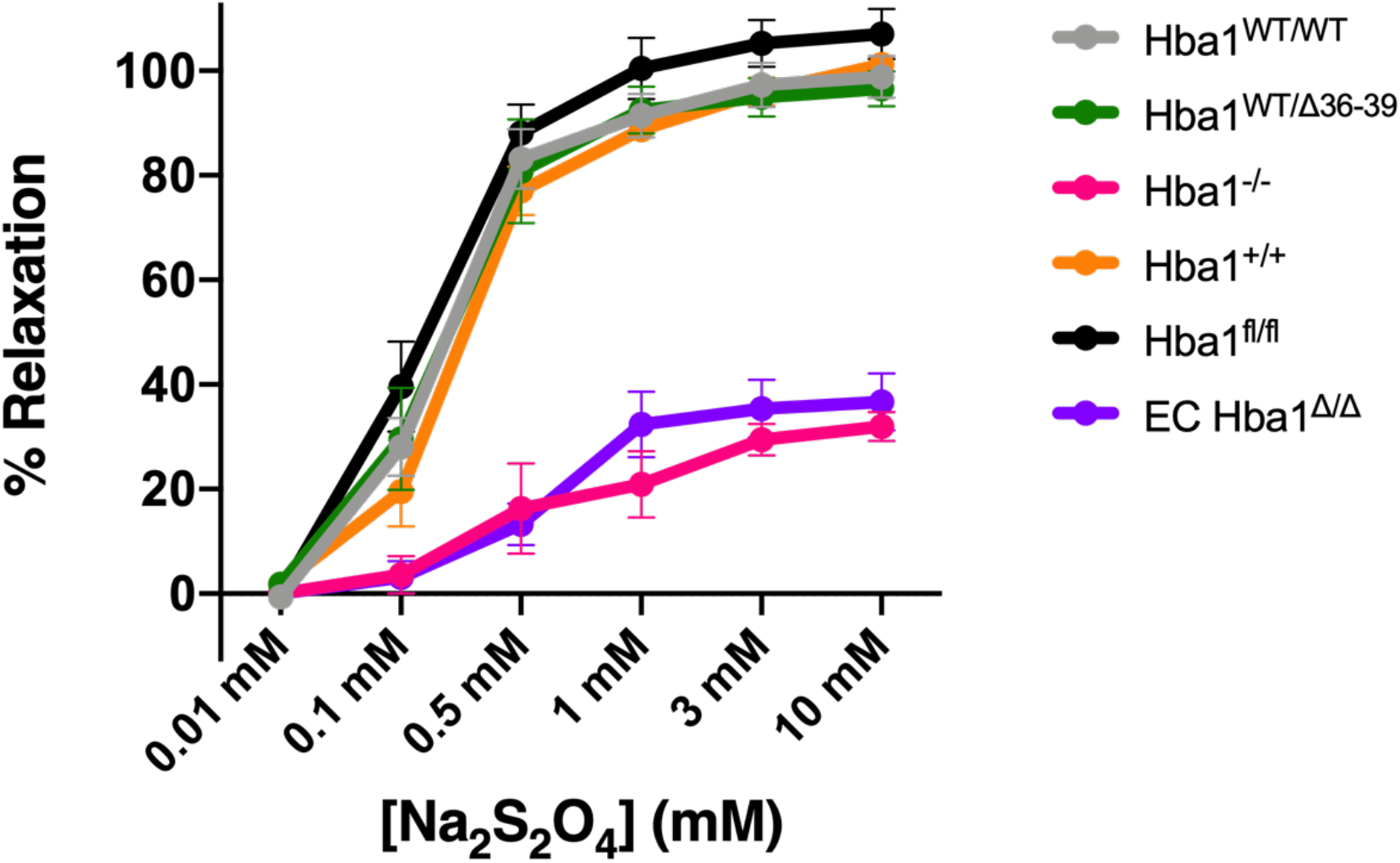
Dose response curve of thoracodorsal arteries from all genetic models to chemical hypoxia. Vasodilation responses to Na_2_S_2_O_4_ are similar between EC-specific (EC *Hba1*^*Δ/Δ*^, purple) and global *Hba1* knockout (*Hba1*^*-/-*^, magenta). Some traces are reproduced from Main Text Figure 6B. Each point on the graph is representative of n > 3 samples per dose and genotype.

**Supplemental Table 1:** Genotypes of P21 pups from five breeding pairs of *Hba1*^*fl/+*^ mice.

| *Hba1<sup>fl/+</sup>* x *Hba1<sup>fl/+</sup>* |  |  |  |  |
| --- | --- | --- | --- | --- |
| Mating Pair | Litter | *Hba1<sup>fl/fl</sup>* | *Hba1<sup>fl/+</sup>* | *Hba1<sup>WT</sup>* |
| 1 | 1 | 1 | 4 | 2 |
|  | 2 | 2 | 4 | 2 |
| 2 | 1 | 3 | 2 | 3 |
|  | 2 | 3 | 3 | 2 |
| 3 | 1 | 2 | 4 | 2 |
|  | 2 | 2 | 3 | 2 |
| 4 | 1 | 1 | 4 | 1 |
|  | 2 | 3 | 1 | 3 |
| 5 | 1 | 1 | 4 | 2 |
|  | 2 | 2 | 5 | 1 |
| Total |  | 20 | 34 | 20 |
| Percentage |  | 27 | 46 | 27 |
| *p-value range: 0.10 - 0.90* |  |  |  |  |

**Supplemental Table 2:** Genotypes of P21 pups from four breeding pairs of *Hba1*^*fl/+*^; *Cdh5-Cre*^*ERT2*^ mice.

| *Hba1<sup>fl/+</sup>; Cdh5-Cre<sup>ERT2</sup>* x *Hba1<sup>fl/+</sup>; Cdh5-Cre<sup>ERT2</sup>* |  |  |  |  |
| --- | --- | --- | --- | --- |
| Mating Pair | Litter | *Hba1<sup>fl/fl</sup>; Cdh5-Cre<sup>ERT2</sup>* | *Hba1<sup>fl/+</sup>; Cdh5-Cre<sup>ERT2</sup>* | *Hba1<sup>+/+</sup>; Cdh5-Cre<sup>ERT2</sup>* |
| 1 | 1 | 1 | 5 | 2 |
|  | 2 | 3 | 2 | 2 |
| 2 | 1 | 2 | 2 | 3 |
|  | 2 | 3 | 3 | 1 |
|  | 3 | 3 | 4 | 0 |
| 3 | 1 | 1 | 4 | 1 |
|  | 2 | 1 | 4 | 3 |
|  | 3 | 4 | 2 | 3 |
| 4 | 1 | 0 | 5 | 2 |
|  | 2 | 1 | 3 | 2 |
| Total |  | 19 | 34 | 19 |
| Percentage |  | 26.4 | 47.2 | 26.4 |
| *p-value range: 0.975 - 0.99* |  |  |  |  |

**Supplemental Table 3:** Genotypes of P21 pups from three breeding pairs of *Hba1*^*fl/+*^; *Sox2-Cre* mice.

| ***Hba1<sup>fl/+</sup>*; *Sox2-Cre* x *Hba1<sup>fl/+</sup>*; *Sox2-Cre*** |  |  |  |  |
| --- | --- | --- | --- | --- |
| **Mating Pair** | **Litter** | ***Hba1<sup>fl/fl</sup>*; *Sox2-Cre*** | ***Hba1<sup>fl/+</sup>*; *Sox2-Cre*** | ***Hba1<sup>+/+</sup>*; *Sox2-Cre*** |
| **1** | 1 | 1 | 5 | 0 |
|  | 2 | 1 | 3 | 1 |
|  | 3 | 1 | 3 | 1 |
|  | 4 | 1 | 3 | 1 |
|  | 5 | 0 | 3 | 3 |
| **2** | 1 | 0 | 3 | 3 |
|  | 2 | 1 | 3 | 2 |
| **3** | 1 | 1 | 3 | 2 |
|  | 2 | 1 | 3 | 2 |
|  | 3 | 1 | 3 | 3 |
| **Total** |  | 8 | 32 | 18 |
| **Percentage** |  | 13.7 | 55.2 | 31.1 |
| ***p*-value range: 0.1 - 0.9** |  |  |  |  |

**Supplemental Table 4:** Genotypes of P21 pups from three breeding pairs of *Hba1*^*+/neo*^ mice.

| ***Hba1<sup>+/-neo</sup>* x *Hba1<sup>+/-neo</sup>*** |  |  |  |  |
| --- | --- | --- | --- | --- |
| **Mating Pair** | **Litter** | ***Hba1<sup>neo/neo</sup>*** | ***Hba1<sup>+/-neo</sup>*** | ***Hba1<sup>+/+</sup>*** |
| **1** | 1 | 0 | 4 | 2 |
|  | 2 | 1 | 3 | 1 |
|  | 3 | 1 | 3 | 2 |
|  | 4 | 1 | 6 | 0 |
| **2** | 1 | 1 | 4 | 2 |
|  | 2 | 0 | 5 | 3 |
|  | 3 | 1 | 3 | 1 |
| **3** | 1 | 0 | 3 | 3 |
|  | 2 | 1 | 2 | 2 |
|  | 3 | 1 | 2 | 3 |
| **Total** |  | 7 | 35 | 19 |
| **Percentage** |  | 11.5 | 57.4 | 31.1 |
| ***p*-value range: 0.025 - 0.05** |  |  |  |  |

**Supplemental Table 5:** Genotypes of P21 pups from three breeding pairs of *Hba1*^*WT/Δ36-39*^ mice.

| *Hba1*<sup>WT/Δ36-39</sup> x *Hba1*<sup>WT/Δ36-39</sup> |  |  |  |  |
| --- | --- | --- | --- | --- |
| Mating Pair | Litter | *Hba1*<sup>Δ36-39/Δ36-39</sup> | *Hba1*<sup>WT/Δ36-39</sup> | *Hba1*<sup>WT/WT</sup> |
| **1** | 1 | 0 | 4 | 4 |
|  | 2 | 0 | 2 | 3 |
|  | 3 | 0 | 2 | 3 |
|  | 4 | 0 | 2 | 5 |
|  | 5 | 0 | 3 | 3 |
| **2** | 1 | 0 | 4 | 4 |
|  | 2 | 0 | 2 | 2 |
|  | 3 | 0 | 2 | 3 |
|  | 4 | 0 | 3 | 3 |
| **3** | 1 | 1 | 4 | 1 |
| **Total** |  | 1 | 28 | 31 |
| **Percentage** |  | 1.67 | 46.67 | 51.67 |
| *p-value* < 0.005 |  |  |  |  |

## References

1. Allen BW, Stamler JS, Piantadosi CA. Hemoglobin, nitric oxide and molecular mechanisms of hypoxic vasodilation. Trends Mol Med 15, 452–460 (2009).

2. Kim-Shapiro DB, Gladwin MT, Patel RP, Hogg N. The reaction between nitrite and hemoglobin: the role of nitrite in hemoglobin-mediated hypoxic vasodilation. J Inorg Biochem 99, 237–246 (2005).

3. Umbrello M, Dyson A, Feelisch M, Singer M. The key role of nitric oxide in hypoxia: hypoxic vasodilation and energy supply-demand matching. Antioxid Redox Signal 19, 1690–1710 (2013).

4. Maher AR, et al. Hypoxic modulation of exogenous nitrite-induced vasodilation in humans. Circulation 117, 670–677 (2008).

5. Tsoukias NM, Kavdia M, Popel AS. A theoretical model of nitric oxide transport in arterioles: frequency-vs. amplitude-dependent control of cGMP formation. Am J Physiol Heart Circ Physiol 286, H1043–1056 (2004).

6. Premont RT, Stamler JS. Essential Role of Hemoglobin betaCys93 in Cardiovascular Physiology. Physiology (Bethesda) 35, 234–243 (2020).

7. Sun CW, et al. Hemoglobin beta93 Cysteine Is Not Required for Export of Nitric Oxide Bioactivity From the Red Blood Cell. Circulation 139, 2654–2663 (2019).

8. Pawloski JR, Hess DT, Stamler JS. Export by red blood cells of nitric oxide bioactivity. Nature 409, 622–626 (2001).

9. Keller TCt, et al. Modulating Vascular Hemodynamics With an Alpha Globin Mimetic Peptide (HbalphaX). Hypertension 68, 1494–1503 (2016).

10. Straub AC, et al. Hemoglobin alpha/eNOS coupling at myoendothelial junctions is required for nitric oxide scavenging during vasoconstriction. Arterioscler Thromb Vasc Biol 34, 2594–2600 (2014).

11. Straub AC, et al. Endothelial cell expression of haemoglobin alpha regulates nitric oxide signalling. Nature 491, 473–477 (2012).

12. Lechauve C, et al. Endothelial cell alpha-globin and its molecular chaperone alpha-hemoglobin-stabilizing protein regulate arteriolar contractility. J Clin Invest 128, 5073–5082 (2018).

13. Ottolini M, et al. Mechanisms underlying selective coupling of endothelial Ca(2+) signals with eNOS vs. IK/SK channels in systemic and pulmonary arteries. J Physiol 598, 3577–3596 (2020).

14. Denton CC, et al. Loss of alpha-globin genes in human subjects is associated with improved nitric oxide-mediated vascular perfusion. Am J Hematol, (2020).

15. Sander JD, Maeder ML, Reyon D, Voytas DF, Joung JK, Dobbs D. ZiFiT (Zinc Finger Targeter): an updated zinc finger engineering tool. Nucleic Acids Res 38, W462–468 (2010).

16. Mali P, et al. RNA-guided human genome engineering via Cas9. Science 339, 823–826 (2013).

17. Hwang WY, et al. Efficient genome editing in zebrafish using a CRISPR-Cas system. Nature biotechnology 31, 227–229 (2013).

18. Mali P, et al. RNA-guided human genome engineering via Cas9. Science 339, 823–826 (2013).

19. Khandros E, Thom CS, D’Souza J, Weiss MJ. Integrated protein quality-control pathways regulate free α-globin in murine β-thalassemia. Blood 119, 5265–5275 (2012).

20. Alter B. Gel electrophoretic separation of globin chains. Progress in clinical and biological research 60, 157–175 (1981).

21. Kong Y, et al. Loss of α-hemoglobin–stabilizing protein impairs erythropoiesis and exacerbates β-thalassemia. The Journal of clinical investigation 114, 1457–1466 (2004).

22. Yu X, et al. An erythroid chaperone that facilitates folding of α-globin subunits for hemoglobin synthesis. The Journal of clinical investigation 117, 1856–1865 (2007).

23. Sorensen S, Rubin E, Polster H, Mohandas N, Schrier S. The role of membrane skeletal-associated alpha-globin in the pathophysiology of beta-thalassemia. Blood 75, 1333–1336 (1990).

24. DeLalio LJ, et al. Constitutive SRC-mediated phosphorylation of pannexin 1 at tyrosine 198 occurs at the plasma membrane. J Biol Chem 294, 6940–6956 (2019).

25. Villalba N, et al. Traumatic brain injury disrupts cerebrovascular tone through endothelial inducible nitric oxide synthase expression and nitric oxide gain of function. J Am Heart Assoc 3, e001474 (2014).

26. Laker RC, et al. Ampk phosphorylation of Ulk1 is required for targeting of mitochondria to lysosomes in exercise-induced mitophagy. Nat Commun 8, (2017).

27. Sangwung P, et al. Regulation of endothelial hemoglobin alpha expression by Kruppel-like factors. Vasc Med 22, 363–369 (2017).

28. Sorensen I, Adams RH, Gossler A. DLL1-mediated Notch activation regulates endothelial identity in mouse fetal arteries. Blood 113, 5680–5688 (2009).

29. Hayashi S, Lewis P, Pevny L, McMahon AP. Efficient gene modulation in mouse epiblast using a Sox2Cre transgenic mouse strain. Mech Develop 119, S97–S101 (2002).

30. Leder A, Daugherty C, Whitney B, Leder P. Mouse zeta- and alpha-globin genes: embryonic survival, alpha-thalassemia, and genetic background effects. Blood 90, 1275–1282 (1997).

31. Paszty C, et al. Lethal alpha-thalassaemia created by gene targeting in mice and its genetic rescue. Nat Genet 11, 33–39 (1995).

32. Casey DP, Madery BD, Curry TB, Eisenach JH, Wilkins BW, Joyner MJ. Nitric oxide contributes to the augmented vasodilatation during hypoxic exercise. J Physiol 588, 373–385 (2010).

33. Lavier J, et al. Supramaximal Intensity Hypoxic Exercise and Vascular Function Assessment in Mice. J Vis Exp, (2019).

34. Richalet JP, Winkler L, Lhuissier FJ. Acute Hypoxia Decreases Systemic Blood Pressure at Exercise in Hypertensive and Normotensive Subjects. Faseb J 31, (2017).

35. Kim-Shapiro DB, Gladwin MT. Mechanisms of nitrite bioactivation. Nitric Oxide 38, 58–68 (2014).

36. Cosby K, et al. Nitrite reduction to nitric oxide by deoxyhemoglobin vasodilates the human circulation. Nat Med 9, 1498–1505 (2003).

37. Shu X, et al. Heterocellular Contact Can Dictate Arterial Function. Circ Res 124, 1473–1481 (2019).

38. Billaud M, Lohman AW, Johnstone SR, Biwer LA, Mutchler S, Isakson BE. Regulation of cellular communication by signaling microdomains in the blood vessel wall. Pharmacol Rev 66, 513–569 (2014).

